# Herbarium genomics reveal signatures of colonization history, lineage turnover, and adaptation during invasion

**DOI:** 10.64898/2026.03.30.715407

**Authors:** Brandon Hendrickson, Courtney M. Patterson, Nevada King, Braden I. Doucet, Fernando Hernandez, Nicholas J. Kooyers

## Abstract

- A key challenge in invasion biology is dissecting the roles of colonization history, admixture, and selection on invasion success. However, it is often difficult to determine the sources and locations of introduction events as well as the timing and magnitude of gene flow and selection using genomics datasets from contemporary samples.
- We disentangle the evolutionary processes acting during the North American invasion of *Trifolium repens* (white clover) using low-coverage whole genome sequencing from 513 herbarium specimens originally collected between 1838-2018.
- Introduction history paralleled European colonization patterns. The earliest collected samples had predominantly Belgian and British ancestry, with increasing French and Spanish contributions over time in the Mid-Atlantic and South. Genomic diversity increased significantly through time likely driven by admixture between independently introduced lineages. In lowest latitude regions, there is a strong shift in dominant ancestry with limited admixture (i.e., lineage turnover). Putatively adaptive loci exhibited protracted shifts in allele frequency during the invasion, consistent with contemporary clines.
- Our study demonstrates the power of herbarium genomics to provide spatiotemporal insight into recent evolutionary history. We find a complex invasion history that includes multiple introductions, lineage turnover, and admixture that increased genetic diversity and likely facilitated adaptation in this now cosmopolitan plant.

## Introduction

Herbarium genomics has begun to provide an unprecedented view into the evolutionary processes that act on ecological timescales. With nearly 400 million herbarium specimens collected within the past four hundred years (Davis, 2023), these collections span the modern era of global change and can provide novel insights on the impact of industrialization, land use change, climate change, and biological invasions on natural plant populations (Heberling *et al*., 2019). Herbarium genomics has increasingly been used to dissect the evolutionary processes occurring during species invasion (Holmes *et al*., 2016; Burbano & Gutaker, 2023; Kim *et al*., 2023). Genomic insights from contemporary samples now suggest that multiple introductions, admixture, and adaptation are all common during the invasion process (Dlugosch *et al*., 2015; Van Boheemen *et al*., 2017; Vallejo-Marín *et al*., 2021; Battlay *et al*., 2025). Yet, genomic analysis of contemporary samples views only the endpoint of the invasion, providing a low-resolution view of evolutionary processes occurring during invasions. Including herbarium specimens collected throughout the invasion increases temporal and spatial resolution to identify key evolutionary events and provides a glimpse into the lineages that failed instead of only the lineages that thrived.

Herbarium genomics has the potential to better uncover the spatial and temporal patterns of colonization history, admixture, and adaptation influencing invasive species. Identifying the number and source of colonizing lineages is an important first step in invasion genomics, yet it is notoriously difficult to reconstruction detailed timelines of colonization history (Dlugosch & Parker, 2008; Bock *et al*., 2015; Cristescu, 2015; Sherpa & Després, 2021). Each potential introduction adds additional genetic variation into the introduced range, simultaneously facilitating adaptation to novel environments as well as obscuring historic bottlenecks and overwriting previous colonization history (Keller & Taylor, 2008; Dlugosch *et al*., 2015; Estoup *et al*., 2016). The genomic profile of herbarium samples may provide a more reliable window into the genetic diversity and ancestry of early colonists (Martin *et al*., 2013; Exposito-Alonso *et al*., 2018; Gutaker *et al*., 2019; Nevill *et al*., 2020). With this additional clarity, connecting invasions to anthropogenic dispersal patterns is likely possible. For instance, genomic time-series are likely necessary to decipher the extent that invasion patterns of non-native species into North America are shaped by the geopolitical and economic legacies of European colonialism.

Herbarium genomics also offers the opportunity to better uncover the timing and extent of admixture and lineage turnover during invasions. Multiple and repeated introductions are more common than previously recognized, especially in horticultural and agricultural species (Kolbe *et al*., 2004; Bossdorf *et al*., 2005; Roman & Darling, 2007; Facon *et al*., 2008; Shirsekar *et al*., 2021; Battlay *et al*., 2025). Admixture between divergent lineages is common during invasions (Roman & Darling, 2007; Dlugosch & Parker, 2008; Prentis *et al*., 2008; Rius & Darling, 2014) and can increase genetic diversity, generate heterosis and produce novel transgressive phenotypes (Verhoeven *et al*., 2011; Rius & Darling, 2014; Bock *et al*., 2015). Although demographic simulations can provide estimates of the extent and timing of admixture, confidence intervals around such estimates are often large and recent admixture events are difficult to detect (Excoffier *et al*., 2013). However, introgression can be more readily detected in the first few generations following admixture when the genomic signature is the strongest. Thus, with dense temporal and spatial sampling, the time and location of contact between different invasion fronts can be readily identified. Such sampling can also be used to detect lineage turnover following contact between invasion fronts; that is, lineages may die out or are replaced by more competitive lineages. Population genomics studies of contemporary samples will have limited evidence of the existence of these relict lineages, but such turnover has already been documented in temporal genomic time-series of other species (Allentoft *et al*., 2024).

Greater insight into the tempo and mode of evolution is also possible with herbarium genomics. Rapid adaptation is now widely documented during biological invasions (Whitney & Gabler, 2008; Colautti & Lau, 2015). Evidence includes the rapid formation of phenotypic and genomic clines across introduced ranges (Kooyers & Olsen, 2012; Colautti & Barrett, 2013; Kooyers & Olsen, 2013; Sotka *et al*., 2018; Gamba *et al*., 2025) as well as fitness advantages for invasive populations relative to native popuations in common garden and reciprocal transplant studies (Colautti & Lau, 2015; Wu & Colautti, 2022). However, these studies provide little insight into the how quickly adaptation occurs following invasion. Rapid selection could occur after introduction or may occur through a more protracted process (Innes *et al*., 2022). Selection could occur across the introduced range or may be more focused on individual regions that have divergent climatic conditions from source populations. Finally, admixture between divergent lineages may or may not play an important role in providing novel genetic diversity for adaptation. Establishing how allele frequencies change through time and space at key candidate loci compared to neutral regions of the genome provides insight into each of these questions.

White clover (*Trifolium repens* L.) is quickly becoming a model species for invasion genomics. White clover is a recently derived allopolyploid (15,000 - 28,000 years ago; 2n = 4x = 32) with disomic inheritance (Griffiths *et al*., 2019). Following hybridization between its parental species, *T. repens* expanded its range across all of Europe and was later domesticated between 1000-1200AD in Spain as a forage and rotational crop (Kjærgaard, 2003). White clover was documented in Britain in the early 1600s and was already naturalized as a feral commensal species across eastern North America by 1749 (Franklin, 1746; Kalm *et al*., 1812; Carrier & Bort, 1916). Population genomics studies on contemporary samples indicate genomic diversity is high within both the native European range and the introduced range in North America. Patterns of genetic diversity suggest that multiple introduction events to North America are likely (Wu *et al*., 2021; Battlay *et al*., 2025). Introduced populations in North America are most closely related to the main colonial European powers in North America (England, France, and Spain) as well as Belgium (Battlay *et al*., 2025), but the spatial and temporal extent that white clover introductions occurred in parallel to European settlement is unclear.

Selection has played a notoriously prominent role in white clover invasions. In common gardens in North America, climate-matched North American white clover populations have higher fitness than climate-matched native European populations (Albano *et al*., 2026). Parallel climate-associated clines have evolved across multiple introduced ranges both in key phenotypes (Kooyers & Olsen, 2012; Battlay *et al*., 2025; Hendrickson *et al*., 2025) and across the genome (Kuo *et al*., 2024). The most prominent example is the rapid evolution of clines in cyanogenesis, the ability to produce hydrogen cyanide after tissue damage (Daday, 1954; Kooyers & Olsen, 2012; Santangelo *et al*., 2022). Cyanogenesis is polymorphic in white clover and its presence or absence involves two Mendelian genes, *Ac/ac* and *Li/li* (Corkill, 1942). The dominant allele for each gene codes for the presence of necessary biochemical components for cyanogenesis: cyanogenic glucosides are encoded by the *Ac* allele and a hydrolyzing enzyme, linamarase, is encoded by the *Li* allele. At the molecular level, recurring gene deletions underlie both recessive alleles (Olsen *et al*., 2007; Olsen *et al*., 2008; Kuo *et al*., 2024). Parallel clines in cyanogenesis have evolved across climatic gradients both in the native and several introduced ranges of white clover (Daday, 1958; Kooyers & Olsen, 2012; Kooyers *et al*., 2014; Innes *et al*., 2022), suggesting strong climate-related selection occurs following introduction. Additionally, parallel climate-associated clines in five large haploblocks form on different continents where white clover was introduced following European colonization. A subset of these haploblocks have distinct fitness effects within common gardens in the native and introduced range that are consistent with the direction of cline evolution (Battlay *et al*., 2025). Given evidence for selection on each of these genomic regions, we would predict strong shifts in allele frequency during the timeframe of white clover invasions.

Here we use an herbarium genomics approach leveraging hundreds of specimens and low coverage whole genome sequencing to create a genomic timeline of the invasion of white clover in North America. Using this dataset in combination with comparisons with the contemporary populations from the native range, we ask: How does the colonization history of white clover compare to European settlement patterns? Do introduced populations exhibit signatures of admixture or lineage turnover in the introduced range? How do patterns of genomic diversity and structure change during invasion? And finally, what is the spatial and temporal basis of adaptation for loci that undergo rapid evolution following introduction? Our results highlights the power of herbarium genomics to address fundamental questions about the interactions of evolutionary processes on ecological timescales.

## Materials and Methods

### Sample Collection

We assembled a dataset of 513 preserved specimens from 24 U.S. herbaria. These specimens were collected between 1838 and 2018 and represent 281 counties across 39 states (Figure 1; Table S1). All specimens included information on the year and county of collection. Detailed sampling procedures are provided in the supplemental methods. We also incorporated contemporary specimens previously collected and sequenced from 18 native-range populations spanning eight countries (N= 693; Table S1)(Santangelo *et al*., 2022; Battlay *et al*., 2025).

**Figure 1:**
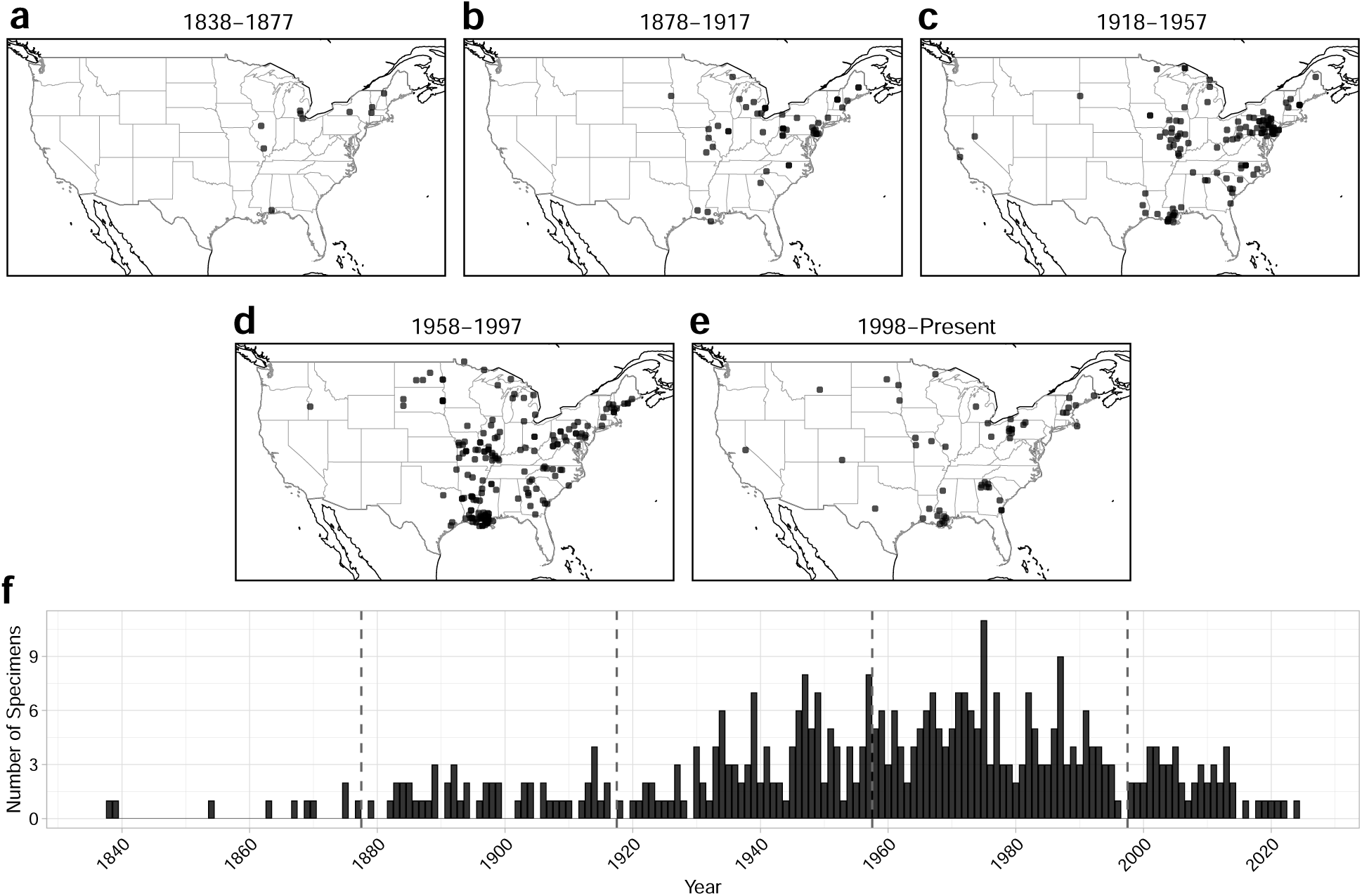
Spatial and temporal distribution of Trifolium repens herbarium collections across the United States from 1838 to the present. Panels show collection localities grouped by time period: (a) 1838–1877, (b) 1878–1917, (c) 1918–1957, (d) 1958–1997, and (e) 1997–present. Each point represents a georeferenced voucher specimen. (f) Histogram showing the frequency of collections by year. Vertical grey dashed lines mark the boundary between time blocks. Herbarium specimens collected before 2000 were prioritized during destructive sampling, and consequently collections after 1998 were not used for downstream analyses.

### DNA Extraction and Library Preparation

Tissue from herbarium samples were stored on silica beads prior to DNA extraction. We extracted DNA using a modified cetyl-trimethylammonium bromide (CTAB) protocol (Doyle & Doyle, 1990; Saeidi *et al*., 2018) with increased the concentrations of CTAB and 2-*β*-mercaptoethanol. We quantified DNA with a Qubit-4 fluorometer (Invitrogen) and diluted all extracts to 3 ng/µL for library preparation. We constructed libraries using a modified dual-index Nextera protocol (Therkildsen & Palumbi, 2017), adjusting the AMPure bead ratio for herbarium material (0.6:1 for contemporary samples; 1:1 for herbarium samples). To monitor contamination, we included negative controls during both DNA extraction and library preparation for herbarium samples to ensure no contemporary DNA entered the workflow. Novogene (Sacramento, CA) sequenced the libraries on HiSeq X (48 samples per lane: lanes 1 & 2) and NovaSeq X Plus (96 samples per lane: lanes 3 through 7) platforms using paired 150-bp reads. We found no batch effects due to differences in sequencing platform in downstream analyese.

### Sequence Processing

A full bioinformatics pipeline can be found on github at github.com/Brandon-Thomas-Hendrickson/HerbariumStructure_WhiteClover. For each contempo-rary sample, we trimmed adapter sequences and poly-G tails using fastp v0.23.4 (Chen *et al*., 2018). We aligned reads to a subgenome and haplotype-resolved *T. repens* reference genome (Santangelo et al. 2023) using bwa mem v0.7.17 (Li & Durbin, 2010) with default settings. We converted SAM files to BAM format, coordinate-sorted them, and removed duplicate reads using samtools v1.18 (Li *et al*., 2009).

Because herbarium specimens likely possess damage and microbial contamination (Bieker *et al*., 2020), we implemented additional steps to reduce the presence of undesirable DNA. First, we trimmed the first and last 5 base pairs of each read using fastp, this step controls for DNA damage which is often concentrated at the termini of degraded herbarium DNA (Bieker *et al*., 2020). Although tools like mapDamage (Ginolhac *et al*., 2011) can assess and correct such damage, we chose not to use them because they require collapsing forward and reverse reads, which significantly reduces usable sequence data, especially when combined with our low-coverage whole-genome sequencing approach. Next, we compiled all complete fungal and bacterial genomes available on NCBI (as of April 24, 2024) into a contamination database and indexed them using Bowtie2 v2.2.4 (Langmead & Salzberg, 2012). We mapped all reads to this database and discarded any reads aligned to the contaminant database. We then aligned the remaining reads using the same pipeline that used for the contemporary samples.

We calculated mapping performance, depth, and coverage statistics for each sample using Qualimap v2.3 (García-Alcalde *et al*., 2012) and Bamtools v2.5.2 (Barnett *et al*., 2011) (Table S1). We removed samples with less than 0.5× genome-wide coverage, which excluded 43 herbarium samples (8.38% of all herbarium samples) and 204 contemporary samples (29.4% of all native range samples). Our final dataset included 470 herbarium samples with a mean coverage of 1.95× (SD = 7.28×) and 489 contemporary samples with a mean coverage of 1.20× (SD = 5.80×) (Table S1).

We calculated genotype likelihoods using ANGSD v0.941-23 (Korneliussen *et al*., 2014) with the Samtools model (-GL 1) and htslib v1.21-27 (Bonfield *et al*., 2021), generating a Beagle file composed of all historic and contemporary samples. We treated the reference allele as the major allele (-doMajorMinor 4) and inferred the minor allele from likelihoods (-doMaf 2). To reduce spurious variant calls, we applied a minor allele frequency (MAF) filter of 0.05 and a SNP p-value threshold of 1e-6. We included only reads with base quality *≥* 20 and mapping quality *≥* 30 (-minQ 20, -minMapQ 30). We filtered sites to retain only four-fold degenerate sites, which we identified using Degenotate v1.3 (Mirchandani *et al*., 2024). Calculation of LD was performed using ngsLD v1.2.0 (Fox *et al*., 2019) with a minimum minor allele frequency of 0.05 and a maximum pairwise distance of 100 kb (–min_maf 0.05, –max_kb_dist 100), and we pruned the resulting LD graph using the prune_ngsLD.py script with a maximum distance of 50 kb and a minimum edge weight of 0.5 (–max_dist 50000, –min_weight 0.5). This filtered dataset contained 172,871 SNPs and served as the basis for all subsequent population structure analyses.

### Reconstructing Colonization History

To infer the geographic origins and colonization pathways of white clover, we integrated multiple complementary ancestry-inference approaches that capture genetic structure at different resolutions and temporal scales. Specifically, we combined principal component–based analyses (PCangsd), supervised classification via random forests, Discriminant Analysis of Principal Components (DAPC), model-based ancestry estimation with NGSadmix, and chloroplast haplotype network reconstruction.

We first characterized broad-scale genetic structure among native-range populations using principal component analysis (PCA) implemented in PCangsd v1.36.2 (Meisner & Albrechtsen, 2018). The resulting covariance matrix was decomposed using the eigen() function in base R v4.4.2 (Team, 2024), and the first ten principal components were retained for downstream analyses. To improve interpretability of population structure and avoid undue influence of extreme samples, we identified and removed 7 outlier herbarium samples based on Mahalanobis distance calculated from PC1 and PC2 scores (outside the 99th percentile; Table S1).

We implemented supervised random forest classification to assign herbarium specimens to their most likely native-range sources. Models were trained to discriminate among native populations using the randomForest package v4.7-1.2 (Liaw & Wiener, 2002). To avoid bias from unequal population sample sizes, we down-sampled all populations to match the smallest reference population (Britain; six individuals). Using these standardized datasets, we performed 1,000 independent PCangsd runs, each based on a random assemblage of six individuals per population. For each iteration, we constructed a training dataset composed solely of native-range samples and a test dataset consisting exclusively of herbarium specimens. Two random forest models were trained per iteration—one using the first ten principal components and one using only the first two—with 5,000 decision trees grown per model. Model performance was evaluated using out-of-bag (OOB) error rates, and the best-performing model across iterations was used to assign herbarium specimens to their most probable source population.

To identify shifts putative source populations over space and time, we applied Discriminant Analysis of Principal Components (DAPC) using the adegenet v2.1.11 package in R (Jombart, 2008). We conducted year-specific DAPC analyses for each year between 1838 and 1997, using subsets of the global covariance matrix that included all eight reference populations and herbarium specimens from the focal year. Each analysis retained 10 principal components (n.pca = 10) and two discriminant functions (n.da = 2), maximizing among-population separation while minimizing within-group variance. Posterior assignment probabilities for herbarium specimens were extracted, normalized, and averaged across samples within each year. This produced a continuous series of inferred source-population contributions, allowing us to track ancestry shifts over time.

We implemented NGSadmix (Skotte *et al*., 2013) to estimate model-based ancestry proportions, which infers individual ancestry while accounting for genotype uncertainty from sequencing data. Ancestry models were evaluated across a range of ancestral population numbers (K = 1–9). For each value of K, five independent runs were performed using distinct random seeds. The optimal K was determined using the Evanno method (Evanno *et al*., 2005) implemented in the CLUMPAK (Kopelman *et al*., 2015).

We constructed haplotype networks from chloroplast genomes using a custom pipeline described in supplemental methods.

### Evidence for Admixture and Lineage Turnover

To track changes in ancestry during the invasion, we estimated a covariance matrix using PCAngsd across all samples and grouped herbarium specimens into 40-year cohorts. We visualized patterns within each cohort to examine temporal shifts in genetic similarity to native populations. We calculated centroids for each native population and 40-year cohort, and then measured Euclidean distances between native and historical centroids. Low distances indicate genetic similarity, whereas high values reflect differentiation. To account for spatial heterogeneity in invasion dynamics, we conducted these analyses separately for northern (>40°N), mid-Atlantic (35–40°N), and southern (<35°N) subsets of the introduced range. These latitudinal regions have distinct evolutionary histories (see Figures S1 and S2) and are therefore analyzed independently in all subsequent analyses.

To directly test for admixture and shifts in ancestral composition, we examined temporal and spatial changes in ancestry proportions inferred with NGSadmix. For each individual, we quantified Max Q – the maximum membership coefficient to any ancestral cluster – using our best fit K (K = 3). Lower values suggesting greater admixture and higher values indicating ancestral purity. We fit both linear and quadratic models of Max Q as a function of year for southern, northern, and mid-Atlantic regions using R::lm(). To detect rare or episodic admixture events not captured by regression models, we identified anomalous individuals relative to a baseline defined by a sliding 30-year average; samples with Max Q values more than 3*σ* below this baseline were classified as outliers suggestive of recent admixture.

Finally, to assess evidence for ancestral lineage turnover, we identify regions where the dominant ancestry flips during the invasion between two informative clusters (Q1 and Q2). We modeled temporal changes of both clusters using R::lm() (i.e., Q1 ∼ Year). Flat slopes in ancestry indicate stability in lineage composition, whereas significant and opposing slopes signal a shift in dominance between ancestral lineages.

### Temporal Patterns of Genetic Diversity in the Introduced Range

We tested for evidence of genetic bottlenecks following introduction by quantifying changes in genetic diversity within the introduced range through time. We calculated individual-level heterozygosity at four-fold degenerate sites. Using both an ancestral consensus genome (Supplemental Methods) and the *T. repens* reference genome (**santangeloHaplotypeResolvedChromosomeLevelAssembly2023a?**), we calculated site allele frequency likelihoods (SAF) for each herbarium sample in ANGSD based on genotype likelihoods (-GL 1, -doSaf 1). To ensure robust estimation, we excluded reads with low mapping or base quality and down-weighted sites with excessive mismatches to reduce overconfidence in variant calls (-C 5). Sample-specific site frequency spectra (SFS) were then inferred from SAF outputs using realSFS under stringent optimization criteria. Model optimization was performed with a maximum of 2000 iterations (-maxiter 2000) and a convergence tolerance of 1e-8 (-tole 1e-8). Individual heterozygosity was calculated as the proportion of the heterozygous genotype likelihood relative to the total likelihood across all three possible genotypes at each site.

We tested for temporal and spatial structure of individual heterozygosity in a linear mixed model framework as a function of collection year, latitude, and their interaction R::lm() (Heterozygosity ∼ Year + Latitude + Year × Latitude). Model adequacy was evaluated using DHARMa package v0.4.7 in R (Hartig, 2016), including formal tests for residual uniformity, dispersion, and outliers. Statistical significance of fixed effects was assessed with ANOVA with a type III sum of square, conducted using the car package v.3.1.3 (Fox & Weisberg).We also modelled change in heterozygosity for samples from the northern, Mid-Atlantic, and southern United States in separate univariate models to better parse regional different in temporal shifts in heterozygosity. Together, these analyses allowed us to track changes in genetic diversity over time.

### Temporal and Spatial Signals of Adaptive Evolution

We examined how allele frequencies shifted over time and space to examine tempo of adaptation during invasions. To explore temporal signals of adaptation for the loci that underlie cyanogenesis, we quantified read abundance at the genomic regions underlying the *Ac/ac* and *Li/li* genes (Olsen *et al*., 2007; Olsen *et al*., 2008; Kuo *et al*., 2024). Read coverage for *Ac* (Chr02_Occ: 9,222,822–9,224,484) and *Li* (Chr04_Pall: 28,511,241–28,515,079) was calculated for each herbarium sample using Samtools (Albano *et al*., 2026). Variation in read counts both reflects presence/absence and copy number variation (Kuo *et al*., 2024). To account for differences in sequencing depth, counts were standardized by the total reads per sample. Standardized read abundance was modeled within linear models with collection year, latitude, and their interaction (Std Read Abundance ∼ Year + Latitude + Year × Latitude) as predictors. Linear models were conducted using the lm() and statistically significance was assessed with ANOVA as above.

Next, we tested whether the frequencies of five putatively adaptive haploblocks have shifted over time in North America (Battlay *et al*., 2025). We generated genotype likelihoods for each haploblock region (Table S2) using ANGSD for each sample, incorporating herbarium samples from the United States and 126 contemporary samples from the GLUE dataset. From the GLUE samples, we selected a subset of individuals representing homozygote reference (AA), heterozygote (Aa), and homozygote alternative (aa) haploblock genotypes, with equal representation of each genotype to avoid bias. We conducted local principal component analyses (PCAs) for each haploblock region using PCangsd (Meisner & Albrechtsen, 2018). We extracted the top 10 genetic principal components via eigen decomposition using the eigen() function in base R. We classified herbarium samples into genotype groups using thresholds along PC1, following Battlay *et al*. (2025), and assigned each sample to one of the three haploblock genotypes (Table S1). To assess whether haploblock genotype frequencies shifted over time and geography, linear regression models were applied with the formula (haploblock genotype frequency ∼ Year + Latitude + Year × Latitude) as above for cyanogenesis loci. This approach enabled the quantification of spatiotemporal trends in haploblock distribution within historical North American white clover populations. We established expectations for cline formation using the common garden studies and contemporary populations genomic analysis presented in Battlay *et al*. (2025).

## Results

### Colonization History Parallels European Settlement Patterns

Colonization history of white clover in North America reflects major contributions from four west European nations: England, France, Spain and Belgium. Introduced populations generally separated from native populations along PC1, with Britain, France and Spain as the most closely related native population to the introduced populations (Figure 2 a). The best fitting random forest model (OOB error rate: 14.58%) finds that the 463 herbarium specimens most closely align with contemporary samples from Britain (280), Spain (143), France (34) and Belgium (6) (Table S3). We find little overlap of herbarium specimens with other sampled native-range populations (Sweden, Greece, Poland, and Germany) in PC space (Figure S3) and none of these populations receive herbarium assignments in the random forest analysis (Table S3). Our chloroplast dataset was not able to provide independent evidence for the above colonization history as most haplotypes were found throughout the key populations in the native range (Figure S4).

**Figure 2:**
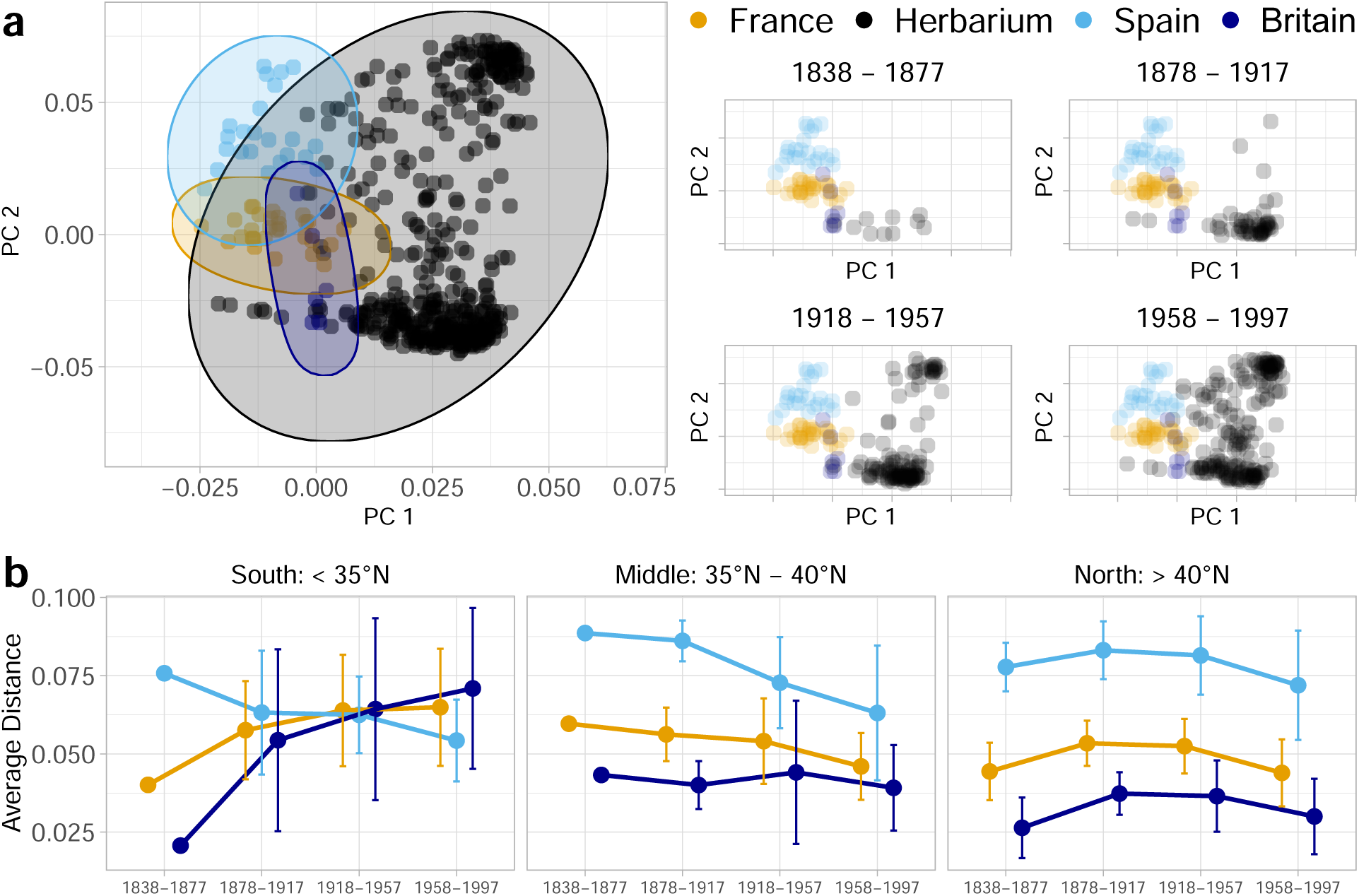
Genetic structure of *Trifolium repens* populations across the native and introduced ranges, and across four 40 year time blocks. (a) Principal component analysis (PCA) of genetic variation. Left panel shows PC1 vs PC2 for all samples (n = 172,871 SNPs) from herbarium specimens (introduced range, black) and contemporary populations from the Britain (dark blue), France (orange), and Spain (light blue). Ellipses represent 95% confidence intervals around country centroids. Right panels show temporal subsets of herbarium specimens across four time periods (1838-1877, 1878-1917, 1918-1957, 1958-1997) plotted with contemporary European populations. Each point represents an individual sample. (b) Temporal changes in genetic similarity between herbarium specimens and European populations. Average Euclidean distances (±SD) between herbarium specimen centroids and contemporary European population centroids across three latitudinal regions: South (<35°N), Mid-Atlantic (35°N-40°N), and North (>40°N). Lines connect temporal means for Britain (dark blue), France (orange), and Spain (light blue) populations. Each point represents the mean distance calculated from PC1-PC2 coordinates, with error bars showing standard deviations across all pairwise comparisons within each time period and region.

Colonization from different source populations occurred at different times and in different areas of the North American range. Subsetting the PCA dataset into four time slices reveals all samples from the early sampling period (1838-1877) are most closely related to Britain and Belgium populations. In the subsequent time slice (1878 -1917), a few individuals begin to appear that are more closely related to France and Spain contemporary samples. The following time bins have even greater contributions from France and Spain. There are clear spatial patterns of introductions associated with different sources – the regions we define as the South, the Mid-Atlantic and the North have distinct evolutionary histories (Figure S1 and S2). A clear latitudinal cline of ancestry (genetic PC2) was established over time, with more recent herbarium collections in the southern United States reflecting Spanish and French ancestry (Figure 2 b) and Britain-derived lineages more common in the northern and Mid-Atlantic regions of the United States. Euclidean distances of genetic PCs show a switch in dominant ancestral composition from British origin to Spanish in the Southern United States occurring after 1950 (Figure 2 b). In contrast, genetic distances between native-range and historic samples does not change in the northern United States – remaining dominated by British lineages. There is also a subtle shift occurring in the Mid-Atlantic region toward Spanish and French ancestry, indicating recent contact between divergent lineages.

Consistent with the principal component analysis, NGSadmix ancestry proportions reveal temporal shifts in ancestry that unfolded in different geographic regions of the United States. We found the most likely number of idealized populations was K=3. All native-range populations possess all three ancestry clusters, but in different proportions, indicating moderate levels of isolation by distance and gene flow in the native range. Britain and Belgium reflect high levels of one ancestry (‘yellow’ or Q1; Figure 3 a), Spain reflect high levels of a second ancestry (‘green’ or Q2), and France is intermediate. The third ancestry cluster (‘purple’ or Q3) is more prominently associated with countries in central and eastern Europe and is not strongly represented in the introduced range. We find that all early herbarium samples across the United States were derived primarily from British populations (Figure 3 b). Overtime, Spanish and French (‘green’) ancestry became more prominent in the southern United States and spread north, however, northern populations remained dominated by British ancestry (‘yellow’). By the 1958-1997 time-block, a clear latitudinal cline in ancestry was established, largely consistant with historical colonization routes of these European countries from the 1500’s – 1800’s.

**Figure 3:**
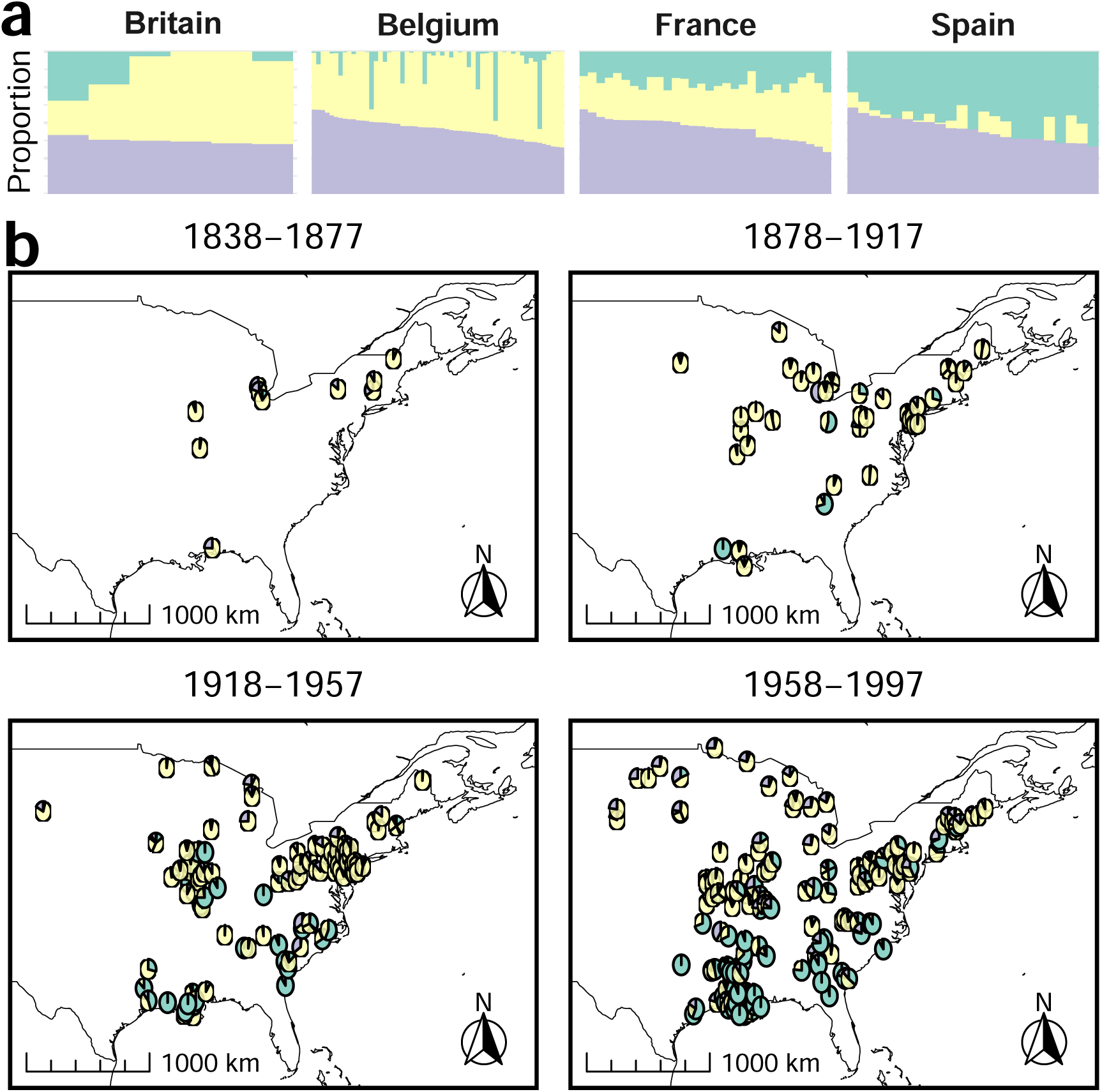
Population genetic structure of *Trifolium repens* within the North American introduced range inferred from NGSadmix analysis (Best K=3). Top panel (a) shows admixture proportions for contemporary populations organized by country (Britain, Belgium, France, Spain). Each vertical bar represents an individual, with colors indicating cluster membership proportions (cluster 1: teal, cluster 2: yellow, cluster 3: purple). (b) Bottom panels show pie charts with ancestry membership coefficients for each North American herbarium samples across four temporal periods. Each pie chart represents an individual and individuals only appear in one of time periods.

### Evidence for both admixture and lineage turnover

There is strong evidence for post-introduction admixture and lineage turnover in different regions of the introduced range. In the Mid-Atlantic region, Max Q significantly declines throughout the 20th century (F_1,98_ = 22.03; *β* = -0.0025; p < 0.0001; Figure 4 b; Table S4), suggesting ongoing admixture between previously isolated clusters. DAPC analyses using 10-year running averages of estimated ancestral proportions are consistent with this shift in the Mid-Atlantic region (p = 0.0067, Figure 4 e; Table S5). Notably, French and Spanish ancestry are increasing (∼0 to 0.16) and while British and Belgian ancestry declined (1 to 0.84) (Table S6) with most change coming relatively recently (i.e., last 80 years). Lineage turnover was more evident within southern North American populations. Britain and Belgium ancestry coefficients declined from ∼1 to 0.088, while France/Spain ancestry coefficients increased from ∼0 to 0.91 over the full time-series (Table S6).

**Figure 4:**
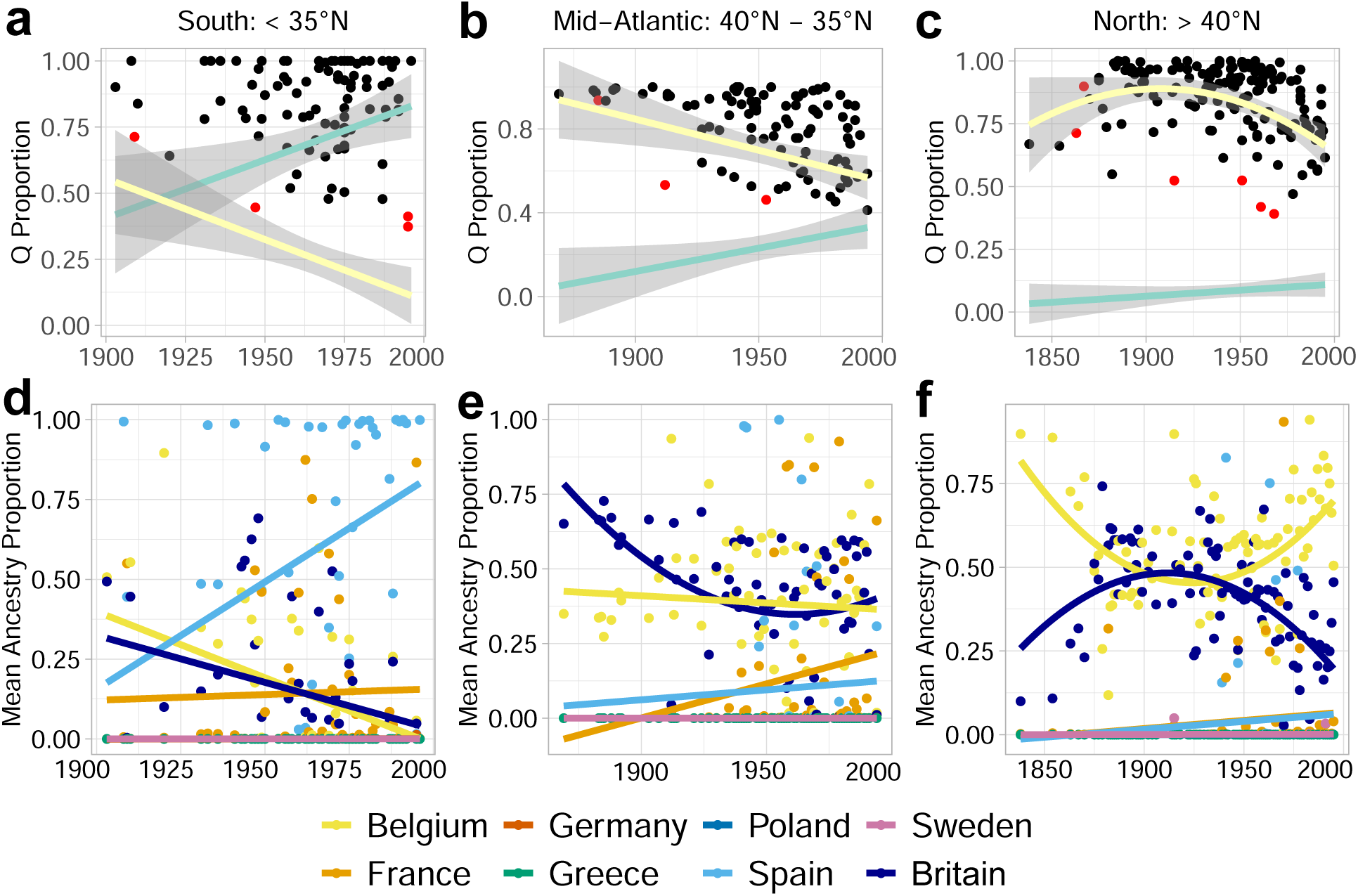
Temporal patterns of genetic cluster membership and ancestry proportions across latitudinal regions in North American herbarium specimens. (a-c) NGSadmix cluster membership coefficients over time (1838-1997) for three latitudinal regions: (a) South (<35°N), (b) Mid-Atlantic (35°N-40°N), and (c) North (>40°N). Each point represents an individual herbarium specimen, with red points indicating statistical outliers (Z-score *≥* 3 from 30-year rolling baseline). Fitted regression lines show temporal trends for cluster 1 (teal) and cluster 2 (yellow). Linear or quadratic models were chosen based on AIC comparison; quadratic relationships shown with polynomial smoothing where appropriate. (d-f) DAPC-derived ancestry proportions over time for the same latitudinal regions: (d) South, (e) Mid-Atlantic, and (f) North. Each point represents mean ancestry proportion per year derived from discriminant analysis of principal components comparing herbarium specimens to eight contemporary European populations. Lines show fitted temporal trends (linear or quadratic based on AIC model selection) for each source country. Analysis based on 10 principal components and 2 discriminant axes, with posterior probabilities normalized across the eight European source populations.

Linear regressions reveal a significantly declining Q1 cluster proportion (F_1,100_ = 9.65; p = 0.0025) and a significantly increasing Q2 cluster proportion (F_1,100_ = 6.91; p = 0.0099), indicating a turnover from British to Spanish ancestral contributions. Linear models of DAPC estimates complement these findings – temporal declines in British ancestry (F_1,46_ = 6.65; p = 0.013) and Belgium ancestry (F_1,46_ = 13.41; p = 0.00064) is synchronous with significant temporal increases in Spanish ancestry (F_1,46_ = 8.062; p = 0.0067). However, unlike in the Mid-Atlantic, Max Q did not exhibit significant changes (F_1,100_ = 0.104; p = 0.75; Figure 4 a). Max Q outliers suggest that admixture was occurring in the Southern United States, but longer-term introgression was limited.

The patterns of admixture and lineage turnover in the Mid-Atlantic and South contrast the relative stability of ancestry over time in the Northern U.S. In the North, there is no significant directional change in Max Q and ancestry coefficients (Figure 4 c). Shifts in ancestry coefficients fluctuate between Belgium and Britain (Figure 4 c; Table S5), a pattern that may or may not have significance since there is little differentiation between these two native regions. A limited number of samples closely related to French and Spanish contemporary samples occur in the north, but there is no progressive rise of either ancestry in this region.

### Genetic diversity increases during the invasion

Neutral genetic diversity increased over the course of the invasion, with greater increases in Mid-Atlantic and southern regions. Individual heterozygosity was highly variable across samples, as expected with heterogeneous sampling across space and time across a species invasion. Individual heterozygosity shifted over time to different extents across the invasive range (Year × Latitude F_1,358_ = 10.31; Year, p = 0.014; Latitude, p = 0.03; Interaction, p = 0.033; Figure 5 a; Table S7). Individual heterozygosity increased through time in the south (F_1,101_ = 10.47; p = 0.0016) and Mid-Atlantic (F_1,98_ = 15.63; p = 0.000145), but only marginally increased in the north (F_1,157_ = 3.218; p = 0.075). Notably, there is a significant correlation between heterozygosity and Max Q, latitude, and their interaction (Table S8). Max Q, a proxy for admixture, is negatively correlated with heterozygosity particularly in the southern (F_1,101_ = 22.39, p = 7.26 x 10-6 ; Figure 5 a) and Mid-Atlantic regions (F_1,98_ = 3.723, p = 0.0566), indicating that greater admixture (i.e., low Max Q) is strongly associated with greater genetic diversity (Table S8). Together, these results are consistent with weak early bottlenecks followed by increasing genetic diversity due to admixture in the southern United States.

**Figure 5:**
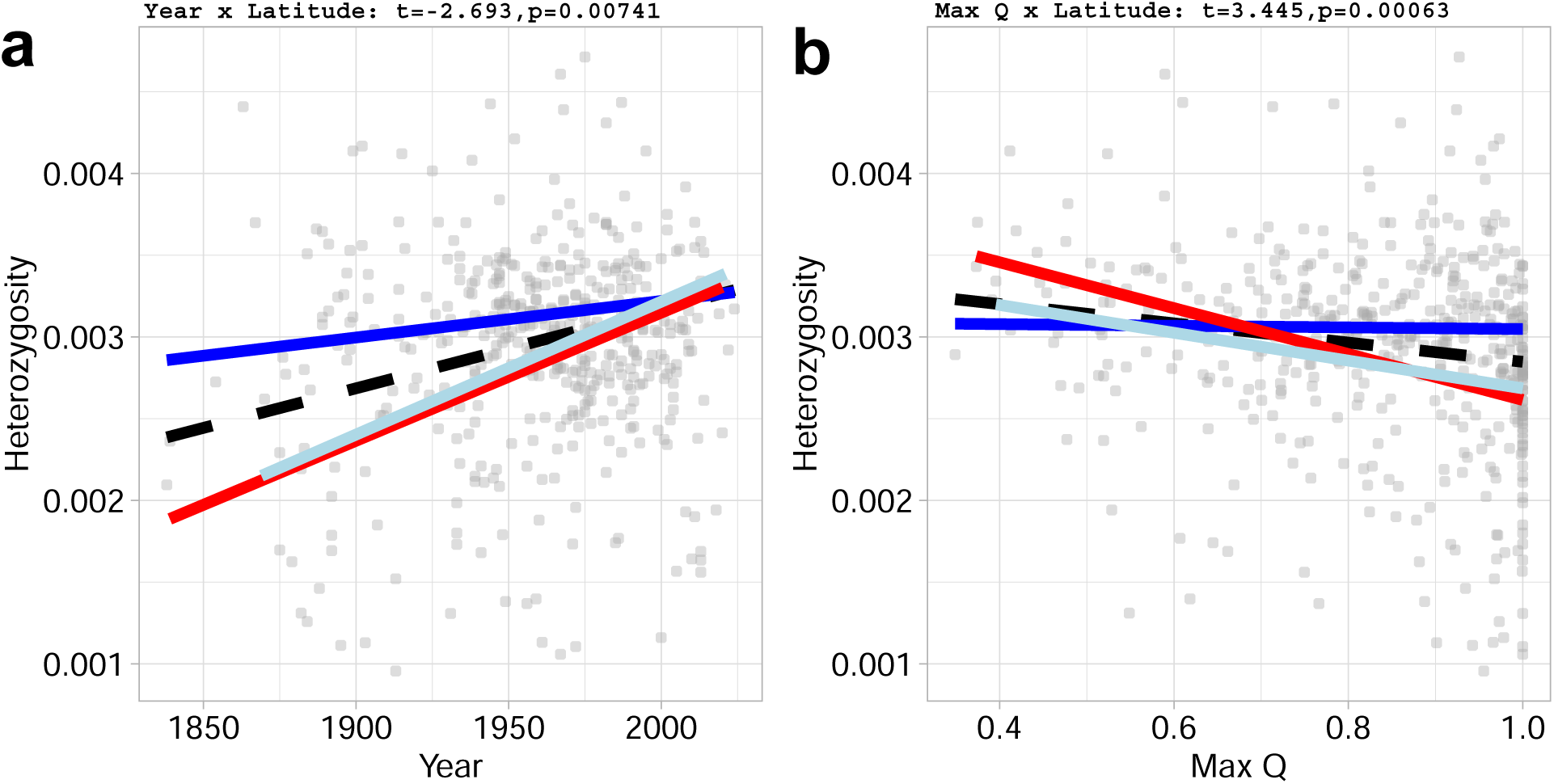
Temporal patterns of genomic diversity in *Trifolium repens*. (a) Individual-level heterozy-gosity patterns over time. Each point represents an individual herbarium specimen (n = 463). Fitted regression lines show predicted heterozygosity trends for northern populations (>40°N, blue solid line), southern populations (<35°N, red solid line), and Mid-Atlantic populations (35°N - 40°N, light blue solid line) and overall trend across all specimens (black dashed line, GLM with Gaussian family: Year × Latitude interaction). (b) The relationship between individual-level heterozygosity with Max Q, proportion of largest ancestral cluster from NGSadmix. All analyses restricted to specimens collected *≤* 1997. Linear models fitted separately for each latitudinal region and combined dataset.

### Candidate genes show temporal signatures of selection across the invasion

We investigated temporal signatures of selection at genomic regions previously identified as rapidly evolving following introduction. Both cyanogenesis-associated loci exhibited significant spatiotemporal patterns in relative read abundance, as evidenced by significant year × latitude interactions (*Ac/ac*: F_1,358_ = 3.814, p = 0.0063; *Li/li*: F_1,358_ = 43.37, p = 0.0018; Table S9). At the *Ac/ac* locus, there were significant effects of year (p = 0.0053) and latitude (p = 0.0068) where read abundance increased throughout the introduction, but were steepest at low latitudes (Figure 6 6). The *Li/li* locus had similar temporal and spatial structure (F_1,358_ = 43.37; year: p = 6.64 × 10^−4^; latitude: p = 0.0029). where read abundance at *Li/li* increased throughout the introduction, and increases were steepest in south. However, in contrast to *Ac/ac*, *Li/li* patterns were driven by a significant temporal increase in southern populations with less temporal change was detected in northern and Mid-Atlantic populations. These results are consistent with the direction and magnitude of latitudinal cyanogenesis clines identified in studies using only contemporary samples (Kooyers & Olsen, 2012; Kuo *et al*., 2024).

**Figure 6:**
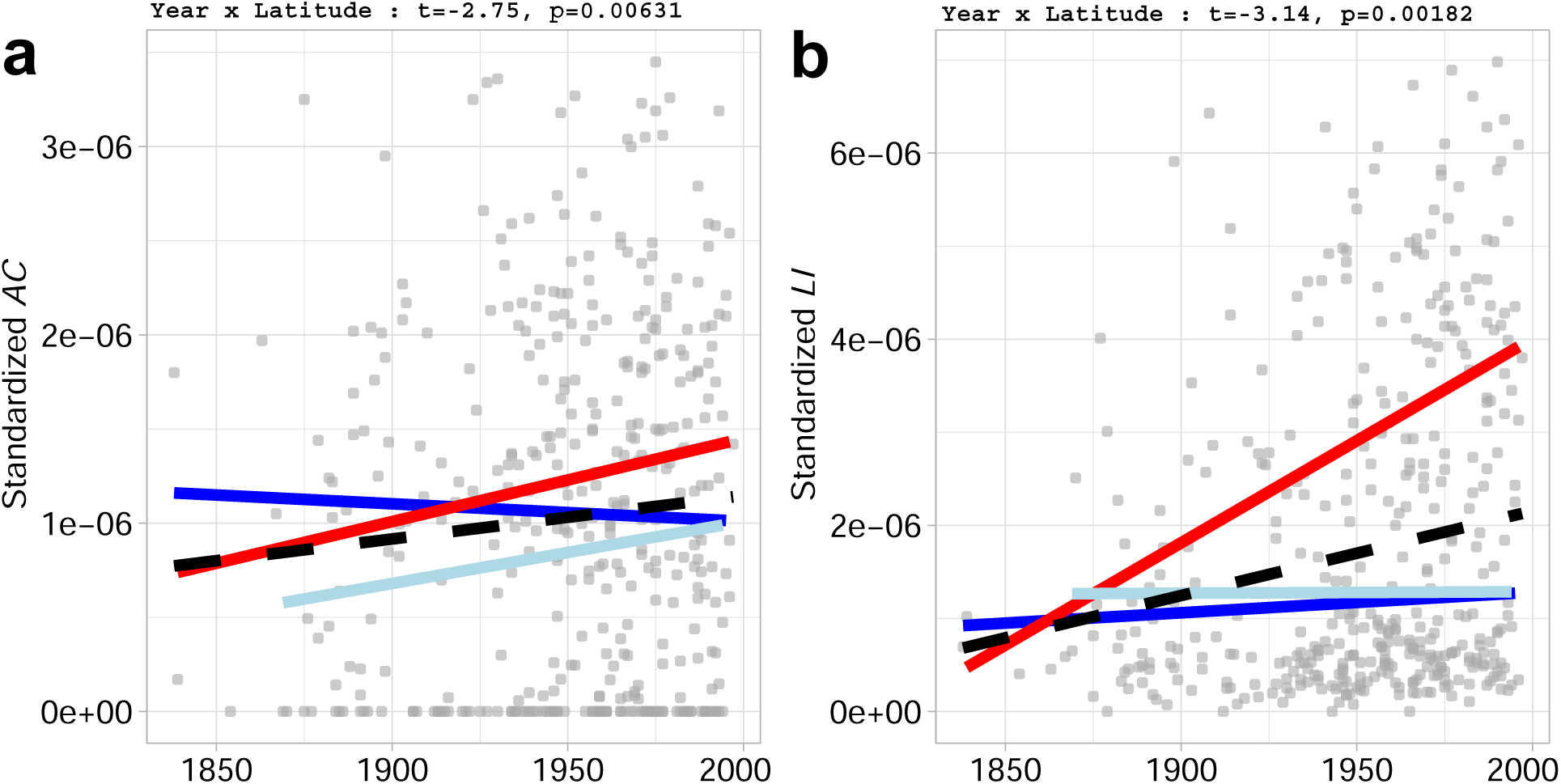
Temporal variation in read number for the genomic regions underlying *Ac/ac* and *Li/li* genes. (a) Temporal trends in standardized read number of Ac (stdAC) across herbarium specimens (1838-1997). Each point represents an individual specimen. Fitted regression lines show predicted trends for northern specimens (>40°N, blue solid), southern specimens (<35°N, red solid), Mid-Atlantic (35°N - 40°N, light blue solid), and overall trend across all specimens (black dashed). (b) Temporal trends in standardized copy number of Li (stdLI) following the same design as panel A.

Unlike the genes involved in cyanogenesis, allele frequencies in the five putatively-adaptive hap-loblocks did not exhibit strong or statistically significant spatiotemporal change in North America (Figure S5; Table S10). Several haploblocks occur at relatively low frequencies in North America (minor haplotype frequencies: hb7a1 = 4.6%, hb7a2 = 26.4%, hb7b = 35.7%, hb9 = 16.1%, hb13 = 11.0%), limiting statistical power to detect modest allele frequency shifts through time. However, we do find some qualitative evidence that haploblock genotypes have shifted in the direction expected from common garden experiments and contemporary population genomic studies. The alternative alleles of both hb7a2 and hb9 – became more frequent in the southern United States whereas the major genotype of hb7b became less frequent overtime in both the southern and northern regions.

## Discussion

Our herbarium genomics approach led to a significant advancement in resolving invasion pathways and deciphering the evolutionary processes acting during the invasion of white clover to North America. Here, we demonstrate that this invasion was not a single, discrete founding event, but an extended process involving multiple introductions from distinct European sources over more than a century. While each introduction involved a substantial pool of initial genetic variation, subsequent contact between distinct introductions led to admixture and increases in genetic variation in the southern and Mid-Atlantic regions of the United States. Notably, contact between distinct introductions also resulted in lineage turnover in the lowest latitude populations. Genomic timeseries at putatively adaptive loci, including the long-studied Mendelian genes underlying cyanogensis, suggest that selection occurred as a protracted process and was heterogenous in intensity across the introduced range. Below we discuss our results in the context of how historical sampling impacts inferences of evolutionary processes acting during invasions and in the context of human colonization history.

### History of Colonization Routes of White Clover

Human activity has long been recognized as the primary driver of biological introductions, with most invasive species closely tied to trade, agriculture, and colonization (Elton, 1958; Crosby, 1986; Lockwood *et al*., 2005). Because Europe was the epicenter of colonial expansion and global trade from the 15th through the early 20th centuries, a disproportionate number of invasive species trace their origins to European source populations (Van Kleunen *et al*., 2015). The genetic structure and spatial distribution of many invasive species are expected to mirror historical settlement pathways, but few contemporary genomics studies have the resolution to test this hypothesis.

We find that the geographic distribution of European white clover lineages in North America aligns with where each respective colonial power settled (Crosby, 1986). Early British and Dutch colonial centers in North America, such as those in modern Mid-Atlantic and New England regions, possess white clover with ancestry similar to native contemporary diversity from those respective countries. In contrast, Spanish and French-associated lineages are observed in the Southern United States, a region that fluctuated between French and Spanish control from 1500-1850. Importantly, we found that independent introductions from the native range unfolded over time, a finding that is only possible by leveraging an historic time-series. Within the temporal and sampling scope of our dataset, we find British lineages arrived first, followed by Spanish and French white clover that subsequently spread throughout the southern United States and, more recently, into the Mid-Atlantic.

Relatively “pure” ancestral lineages—defined as samples with a Max Q of 1—are present within parts of the introduced range of white clover, whereas no native-range samples exhibit this pattern. The patterns are consistent with predictions from invasion genetics and range expansion theory, where founder events and expansion dynamics can allow particular source lineages to establish regionally dominant genetic signatures (Excoffier *et al*., 2009; Estoup & Guillemaud, 2010). Similarly, admixture analyses tend to identify bottlenecked populations as pure clusters (Lawson *et al*., 2018). In the context of biological invasions, such patterns arise because introduced populations typically represent a subset of the genetic diversity present in the native range, often capturing discrete lineages that then expand rapidly into new regions. As a result, early colonists can leave a lasting genetic imprint, producing spatially distinct sectors of high ancestral purity, even in the face of later gene flow. Our temporal herbarium dataset provides direct empirical support for this process.

Several caveats limit inference about the earliest phases of colonization. Our sampling is sparse in key regions of early Spanish settlement, including Florida and southern Texas, limiting our ability to determine when and where Spanish-associated lineages first established in North America. We also cannot easily tease apart French ancestry from Spanish ancestry as native French populations appear as intermediate between Britain and Spain in both PCA and NGSadmix analyses. Notably, we do not see longitudinal relationships with PC2 that would be indicative of early French colonization west of the Appalachian Mountains or in Canada. A final caveat is that our earliest herbarium specimens postdate initial European colonization by centuries; although white clover was described as naturalized by the 18th century (e.g., Franklin (1746)), we lack genomic data from those earliest introductions. Thus, while the observed spatial genetic patterns are consistent with documented settlement routes and trade networks from the 1500s–1800s (Crosby, 1986), they should be interpreted as reflecting subsequent population spread and admixture rather than the precise genetic composition of initial founding populations.

That species invasions closely parallel human migration and colonization is not unique to our study. Parallel spatial patterns between human migration and species invasions have been observed in several other systems: *Arabidopsis thaliana* shows multiple introductions from distinct European origins into North America (Durvasula *et al*., 2017; Exposito-Alonso *et al*., 2018) and *Plantago major* reflects human-mediated dispersal along colonial routes (Ahlstrand *et al*., 2022). Similarly, animal systems, such as *Mus musculus* (Suzuki *et al*., 2013) and *Sturnus vulgaris* (Bodt *et al*., 2020), reveal genetic imprints of colonial history, underscoring that human migration and trade are deeply intertwined with the global movement of species. However, our results suggest a strong temporal lag follows human migration, supporting the hypothesis that invasion ‘debts’ accumulate over time (Essl *et al*., 2011; Rouget *et al*., 2016).

### History of Admixture and Population Turnover

Admixture betwen multiple introductions is thought to be common and conducive to invasion success (Dlugosch & Parker, 2008; Dlugosch *et al*., 2015), however, it is difficult to assess the extent, location and timing of admixture from contemporary population genomic data alone. Our analyses of contemporary and historical white clover provide clear empirical evidence that post-introduction admixture occurs to varying levels across an introduced range. The highest levels of admixture occurs in the Mid-Atlantic and Southern regions, with limited evidence in the Northern region. Interestingly, there are individuals (outliers) with high Spanish and French ancestry that occur in the North, but there is limited introgression. This suggests that there may be selection genome-wide against this ancestry. Greater evidence for selection on the loci and phenotypes associated with this ancestry needs to come from common garden experiments.

There also appears to be more limited introgression within southern region populations (Figure 4) despite clear contact between independently introduced lineages and a distinct shift from British to Spanish ancestry. Such replacement of established lineages with new lineages, i.e., lineage turnover, during the timespan of an invasion has rarely been documented. For instance, there is limited evidence for lineage turnover during the colonization of Europe by *Arabidopsis thaliana* (Gamba *et al*., 2025), but lineage turnover does occur across Europe during the invasion of *Ambrosia artemisiifolia* (Bieker *et al*., 2022). In the southern United States, we find evidence of lineage turnover from British to Spanish ancestry without strong signatures of admixture. We hypothesize that lineage turnover in white clover was driven by environmental filtering, where lineages with Spanish ancestry may have been better adapted to the hot, high-evapotranspiration climate in the southern United States than more British-derived populations. We suggest that lineage turnover is most likely in widespread species that exhibit regional adaptation across their native range, as observed in white clover (Santangelo *et al*., 2022; Innes *et al*., 2022; Albano *et al*., 2026). If multiple different source populations with different environmental tolerances are then introduced to a new area, there will likely be regional establishment of lineages that match local conditions. It is important to note that lineage turnover did not occur in Mid-Atlantic populations. We suggest that neither lineage was better adapted to this region of the introduced region, leading to more time for gene flow. Together, these results highlight the diversity of post-introduction dynamics within an introduced range and demonstrate that distinct evolutionary processes can shape regional patterns of ancestry.

### History of Neutral and Adaptive Loci in White Clover

Early introductions are expected to reduce genetic diversity through founder events and strong genetic drift during colonization (Nei *et al*., 1975; Dlugosch & Parker, 2008). However, herbarium genomes reveal that diversity in *Trifolium repens* increased through time in regions where independently introduced lineages came into contact. Mid-Atlantic populations exhibit clear increases in heterozygosity coinciding with the emergence of mixed Britain–Spain ancestry. Similar patterns have been documented in other invasive systems where secondary contact among introductions restored or elevated genetic diversity (Kolbe *et al*., 2004; Lavergne & Molofsky, 2007; Roman & Darling, 2007; Rius & Darling, 2014). Our temporal dataset directly captures this process: genomic diversity rises following admixture events, demonstrating how multiple introductions can replenish genetic variation during invasion.

Increases in genomic diversity due to admixture could strongly impact adaptive capacity. Southern populations exhibit substantial increases in genome-wide heterozygosity over the temporal series, coinciding with rapid shifts at putatively adaptive loci. It is tempting to suggest that admixture increased the pool of genetic diversity and facilitated adaptive evolution. This finding would be consistent with observations in other invasive systems, including lizards (Kolbe *et al*., 2004; Kolbe *et al*., 2008), aquatic organisms (Roman & Darling, 2007), and plants (Dlugosch & Parker, 2008; Hodgins *et al*., 2018), where secondary contact among independently introduced lineages increased genetic diversity and facilitated adaptive evolution. However, an equally likely scenario is that newly arriving lineages had greater fitness than established lineages, and any introgression between the lineages consisted of neutral variants. Further work is necessary to parse haplotypes at individual loci under selection as well as determine whether adaptive combinations of haplotypes at different loci stem from admixture between distinct lineages.

Our results do reveal that adaptive evolution is has occurred and is ongoing in invasive *Trifolium repens* populations across North America. Specifically, we find selection has increased the frequency of the *Ac* and *Li* alleles (or their copy number) over time, particularly in southern populations. This result matches the prediction from contemporary distributions of cyanogenesis across the native and introduced regions, where low latitude populations have greater frequencies of cyanogenesis (Daday, 1954; Kooyers & Olsen, 2012; Kooyers & Olsen, 2013; Santangelo *et al*., 2022). Notably, we had very limited sampling western U.S. which prevents our study from identifying signatures of selection on the *Ac/ac* locus that may arise from other functions, such as drought resistance (Kooyers *et al*., 2014). Two key observations come the herbarium genomics time series. The evolution of cyanogenesis clines appears protracted in nature, where clines at the *Ac/ac* and *Li/li* have steepened gradually through time rather than rapidly following initial introduction. These results are consistent with a previous resurvey of Daday’s populations across continents, demonstrating that cyanogenesis clines have become steeper and better resemble the cyanogenesis cline in the native region over the last 60 years (Innes *et al*., 2022). Second, the patterns of selection are very similar between both loci and involve far stronger selection in southern populations compared to northern populations. Thus, selection is regional and coincides with lineage turnover in the southern U.S., although the intensity of selection at *Ac/ac* and *Li/li* is far stronger than at neutral loci.

We find less evidence for selection on large haploblocks that have been implicated in adaptation following introductions across the world (Battlay *et al*., 2025). The predicted shifts across space and time are less clear for these genomic regions as some of these haploblocks do not have clear latitudinal patterns within population genomics studies using contemporary data or do not have differences in fitness between low and high latitude common gardens. However, we expected the reference allele to always be favored in the introduced range for hB13, and find only a small insignificant increase through our temporal dataset. We also can predict that the alternative allele for hb7a2 and hb9 should increase more quickly in high latitude populations than low latitude populations, but our dataset suggests both loci have modest range wide increases through time in the alternative alleles. Because alternative alleles are often at relatively low frequency and we have a limited number of herbarium specimens, we likely do not have sufficient power to assess latitude x time effects on genotype frequencies. We note that haploblocks are also polymorphic in most introduced populations across the globe, suggesting that neither allele has been swept to fixation. Together, the absence of strong temporal shifts in haploblock frequencies is largely consistent with expectations from contemporary data, reflecting change through time consistent with weak selection that is spatially or temporally variable, rather than the strong directional change observed in the cyanogenesis loci.

Our results align with a growing body of herbarium genomic studies demonstrating rapid evolutionary change following invasion. Temporal genomic sampling of herbarium specimens in other systems has revealed allele frequency shifts during invasions, including selection on standing genetic variation and climate-associated loci (Martin *et al*., 2013; Van Boheemen *et al*., 2017; Hodgins *et al*., 2018). Similarly, historical and contemporary genomic data has been used to identify adaptive responses to anthropogenic pressures such as agricultural intensification and climate change (Kreiner *et al*., 2018, 2022). While this study has focused on the tempo and mode of selection acting on previously identified candidate genes, a clear next step is to identify signatures of selection genome-wide and assess temporal and spatial generalities across a less-biased set of loci.

### Conclusions

The invasion of *T. repens* into North America emerges as a prolonged, dynamic process rather than a single historical event. Multiple introductions from distinct parts of the native range, temporal turnover in dominant ancestries, admixture-fueled increases in diversity, and adaptation to local environments have all contributed to the species’ success. By leveraging herbarium specimens to generate a century-spanning genomic time series, we find lineages that were once regionally dominant but are now rare, absent, or admixed, highlighting that contemporary genetic structure represents a filtered subset of historical diversity. By providing fine-scale temporal resolution, historical specimens allow us to reconstruct when and where admixture occurred, identify replaced ancestral lineages, track changes in genome-wide diversity, and identify the tempo of adaptation. This integrative approach parallels other successful reconstructions of invasion history (Estoup & Guillemaud, 2010; Cristescu, 2015) and can be applied broadly to reconstruct the genomic histories of other widespread invasive species, providing critical insights into the interplay between demography, selection, and genomic architecture during speices invasions.

## Acknowledgements

We gratefully acknowledge the herbaria and curatorial staff that generously provided access to specimens and sampling permission: the University of Louisiana, Lafayette (LAF), University of Georgia Herbarium (UGA), the Shirley C. Tucker Herbarium at Louisiana State University (LSU), the Missouri Botanical Garden Herbarium (MO), the Academy of Natural Sciences of Drexel University Herbarium (PH), the Vanderbilt University Herbarium (VDB), the University of Illinois Herbarium (ILL), the Carnegie Museum Herbarium (CM), the Black Hills State University Herbarium (BHSU), the University of Michigan Herbarium (MICH), the University of North Carolina at Chapel Hill Herbarium (NCU), the West Virginia University Herbarium (WVA), the University of Southern Mississippi Herbarium (USMS), the University of Mississippi Herbarium (MISS), the Botanical Research Institute of Texas Herbarium (BRIT), the R.L. McGregor Herbarium at the University of Kansas (KANU), University of Arkansas (UARK), the Albion R. Hodgdon Herbarium at the University of New Hampshire (NHA), the C.A. Taylor Herbarium at South Dakota State University (SDSU), and the North Dakota State University Herbarium (NDSU). Their stewardship of historical collections made this study possible. Marc Johnson, James Santangelo, Simon Innes, and Lucas Albano generated genomic data published in past studies and provided support for our work. Nevada King and Aurora Gaspard assisted with specimen collection and molecular work. We are grateful to Matthew Austin, James Beck, Erin Sigel, Craig Barrett, Jonas Mendez-Reneau, Maribeth Latvis, and Donna Hinrichs for assistance with herbarium sampling and curation. This work benefited from computing resources and technical support provided by the Louisiana Optical Network Infrastructure (LONI). We also acknowledge the CPING consortium for collaborative support and the NSF EPSCoR-funded CPING program for financial support of consortium activities. This work was funded by a National Science Foundation grant to NJK (OIA-1920858).

## Data accessibility

All data and code used in this study are available on github (https://github.com/Brandon-Thomas-Hendrickson/HerbariumStructure_WhiteClover) and sequences are deposited in the NCBI Sequence Read Archive (SRA) under BioProject PRJNAxxxxxx.

