## Supplemental Information for "Herbarium genomics reveal signatures of colonization history, lineage turnover, and adaptation during invasion"

Article acceptance date: TBD

The following Supporting Information is available for this article:

### Sample Selection and Data Curation

Herbarium specimens were selected based on the following criteria: (i) the specimen contained 10–20 trifoliate leaves, (ii) the specimen included georeferencing information (latitude/longitude and/or county of collection), and (iii) the date of collection was recorded. To conserve herbarium samples, limit sequencing effort, and reduce spatial and temporal sampling redundancy, only one specimen per county per year was sampled; additional specimens collected within the same county and year were excluded from sampling. The catalog number and herbarium-specific barcode for each specimen were recorded to allow cross-referencing with the SERNEC database and the corresponding herbarium records.

Depending on specimen size, 1–5 trifoliate leaves were removed for DNA extraction. Leaves were placed in coin envelopes and sealed in plastic bags containing silica drying beads for desiccation. All sampled vouchers were modified to include a label indicating the current project title, collector name, and date of destructive sampling. When permitted by the herbarium curator, a small arrow was drawn on the voucher sheet to indicate the leaf that had been removed.

### Chloroplast Networks

We constructed haplotype networks from chloroplast genomes using a custom pipeline. Samples with mean chloroplast genome coverage below 10× were excluded (N = 8), and those with extremely high coverage (>300×) were down sampled to 300x (N = 275 samples). Genotype likelihoods for each site across all chloroplast genomes were estimated using bcftools v1.21 (Danecek *et al.*, 2021) with the mpileup command. Variant calling was then performed using the call -mv function, identifying both SNPs and indels. Resulting variant calls were filtered to retain sites with a minimum mapping quality of 30 and a minimum depth of 10. Additional filtering was performed using VCFtools v0.1.16 (Danecek *et al.*, 2011) to retain only biallelic variants and those with a minor allele frequency (MAF) ≥ 0.01. Missing sites were imputed using Beagle v5.4 (Browning *et al.*, 2018) on normalized variant calls generated with the norm option in bcftools. After filtering, 307 variant chloroplast sites were retained. The final imputed VCF file was converted to PHYLIP format using a custom Python script (Ortiz, 2019) and subsequently converted to NEXUS format using the ape package v5.8 (Paradis & Schliep, 2019). Haplotypes were inferred using the haplotype function from the pegas package v1.3 (Paradis, 2010). To simplify the networks for interpretation, haplotypes represented by only one individual were filtered out. TCS haplotype networks were then constructed using the PopART GUI v1.7.2 (Leigh & Bryant, 2015).

### North and South Distinctions

We fit a log-linear regression model with PC2 as the response variable and the natural log of absolute latitude as the predictor to assess the relationship between ancestry and geographic position . To identify potential geographic transitions in ancestry, we conducted changepoint analysis using the changepoint v.2.3 package in R (Killick, 2011) to estimate the latitude at which the shift between predominantly northern and southern genetic signatures occurs. The log-linear regression revealed a rapid decline in PC2 at higher latitudes ($F1,415 = 358$, $R2 = 0.4631$, $p < 0.0001$; Supplemental Figure 2B), with a breakpoint estimated at 37.77°N.

To identify potential contact zones between geographic clusters, we examined patterns of genetic diversity across latitude using a sliding window approach. Variance in principal components (PCs), used as a proxy for genetic heterogeneity, was calculated within 5° latitude windows advanced in 1° increments from 27°N to 50°N. Within each window, we also quantified the number of individuals with intermediate ancestry proportions (0.1 < Cluster 1 proportion from NGSadmix < 0.9), which served as a proxy for hybridization rates. Subsequent analyses were conducted separately for specimens assigned to each geographic region and any hybrid zone(s) identified. The latitude window spanning 35–40°N exhibited elevated genetic variance and a higher proportion of ancestrally mixed individuals relative to other regions (Supplemental Figure 2A), consistent with a hybrid zone. Based on these patterns, we defined three geographic regions for subsequent analyses: samples collected below 35°N ("south"), between 35–40°N ("mid-Atlantic"), and above 40°N ("north").

**Construction of Ancestral Census Sequence**

An ancestral reference sequence was required to estimate individual heterozygosity from the site frequency spectrum (SFS). Whole-genome sequence data from *Trifolium pratense* (SRA: ERR3481780) and *Trifolium occidentalis* (SRA: SRR8593471) were therefore aligned to the *T. repens* reference genome (Santangelo *et al.*, 2023). These alignments were used to construct an ancestral consensus genome using ANGSD with the -doFasta 2 option. During this step, low-quality reads and duplicates were filtered (-minMapQ 25, -minQ 20, -remove_bads 1, -uniqueOnly 1), and base counts were recorded using the -doCounts 1 option.

**Fig. S1** Ancestry-based population structure differentiates native and introduced *Trifolium repens* populations along primary axes of genetic variation. (a) Principal Component 1 (PC1) variation between native European populations and introduced North American populations. Boxplots show median, quartiles, and range of PC1 values for specimens from Europe (native range, black) and North America (introduced range, grey). (b) Principal Component 2 (PC2) variation between ranges following the same design as panel A. PC1 captures the primary axis of differentiation between European source populations and North American herbarium specimens. (c) Relationship between ancestral proportion (Cluster 1 from NGSadmix K=3 analysis) and PC1. Each point represents an individual specimen. Red line shows fitted linear regression with confidence intervals depicted as grey shading. Ancestral proportion of cluster 1 is used as the predictor of the regression. (d) Relationship between ancestral proportion (Cluster 1) and PC2. Points represent individual specimens. This relationship demonstrates the temporal and ancestry-based structure captured by PC2.

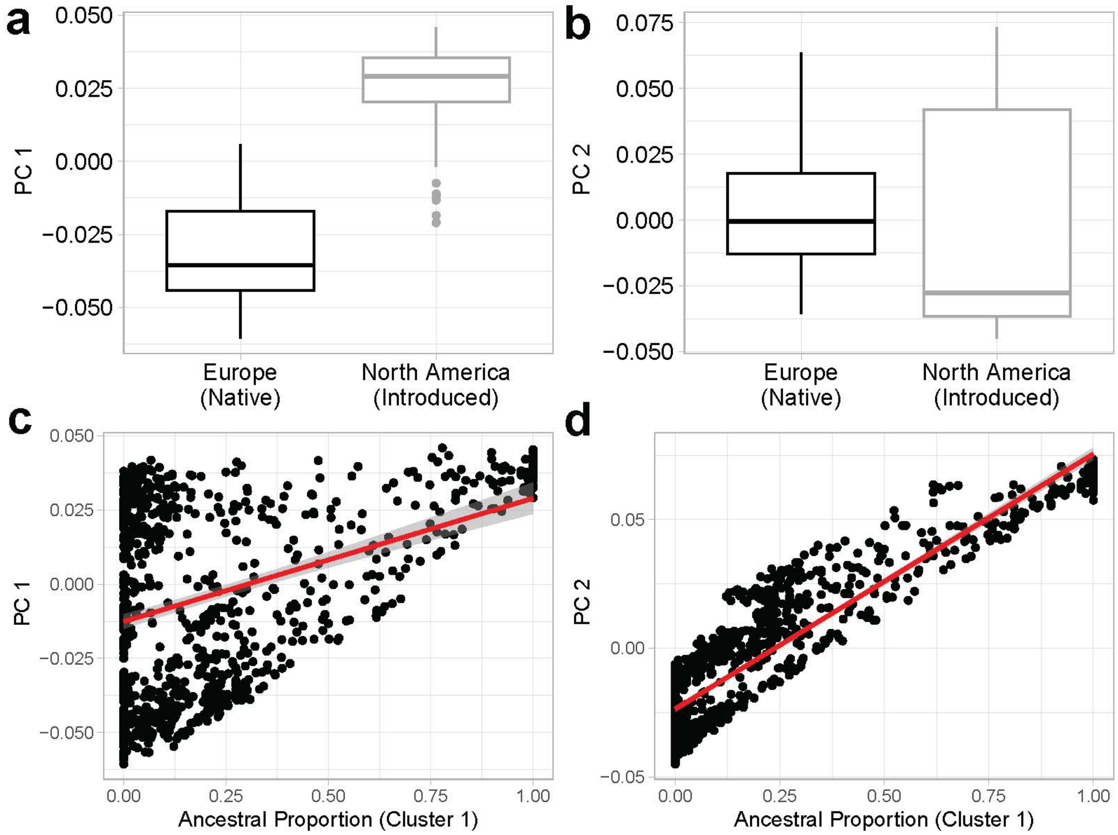

**Fig. S2** Latitudinal gradients in genetic admixture reveal north-south genetic clustering and transition zone in introduced *Trifolium repens* populations. (a) Sliding window analysis of genetic diversity patterns across latitude. Red line shows the count of specimens with intermediate ancestral proportions (0.1 < Cluster 1 proportion < 0.9) within 5-degree latitude windows, indicating zones of genetic admixture. Black line shows PC2 variance within the same latitude windows (scaled to secondary y-axis), representing genetic heterogeneity. Windows advance by 1-degree increments from 27°N to 50°N. Peak counts of admixed individuals occur in the mid-latitudes (35°N-40°N), corresponding with elevated PC2 variance. (b) Latitudinal distribution of herbarium specimens showing regional classification boundaries. Points represent individual specimens colored by region: North (>40°N, blue), South (<35°N, red), and Mid-Atlantic (35°N-40°N, light blue). Vertical dashed line indicates the optimal breakpoint identified through changepoint analysis, marking the transition between predominantly northern and southern genetic signatures.

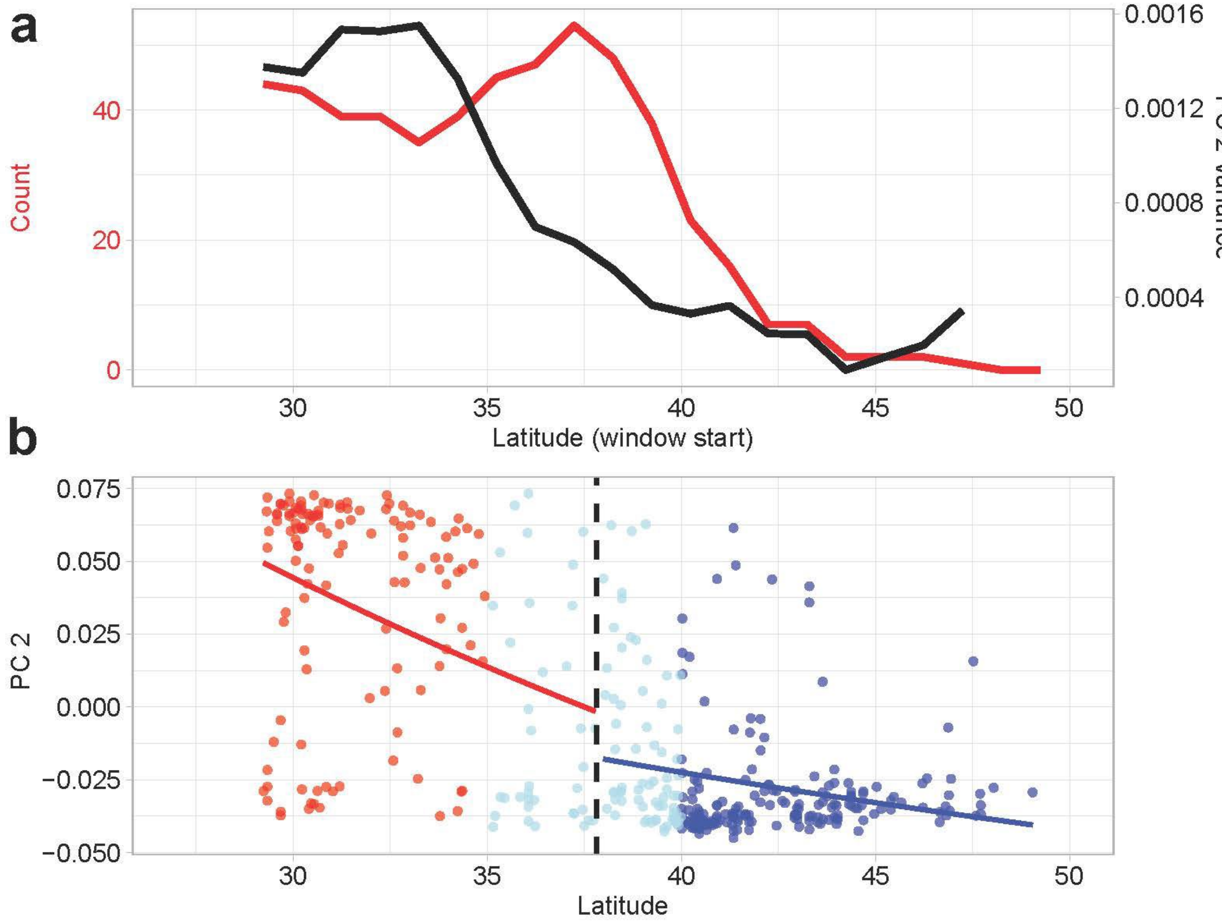

**Fig. S3** Genetic structure of *Trifolium repens* populations across the native (all populations) and introduced ranges, and across four 40-year time blocks. (a) Principal component analysis (PCA) of genetic variation. Left panel shows PC1 vs PC2 for all samples (n = 172,871 SNPs) from herbarium specimens (introduced range, black) and contemporary populations from the Britain, Belgium, Germany, Poland, Sweden, France, Greece, and Spain. Ellipses represent 95% confidence intervals around country centroids. Right panels show temporal subsets of herbarium specimens across four time periods (1838-1877, 1878-1917, 1918-1957, 1958-1997) plotted with contemporary European populations. Each point represents an individual sample. (b) Temporal changes in genetic similarity between herbarium specimens and European populations. Average Euclidean distances (±SD) between herbarium specimen centroids and contemporary European population centroids across three latitudinal regions: South (<35°N), Mid-Atlantic (35°N-40°N), and North (>40°N). Each point represents the mean distance calculated from PC1-PC2 coordinates, with error bars showing standard deviations across all pairwise comparisons within each time period and region.

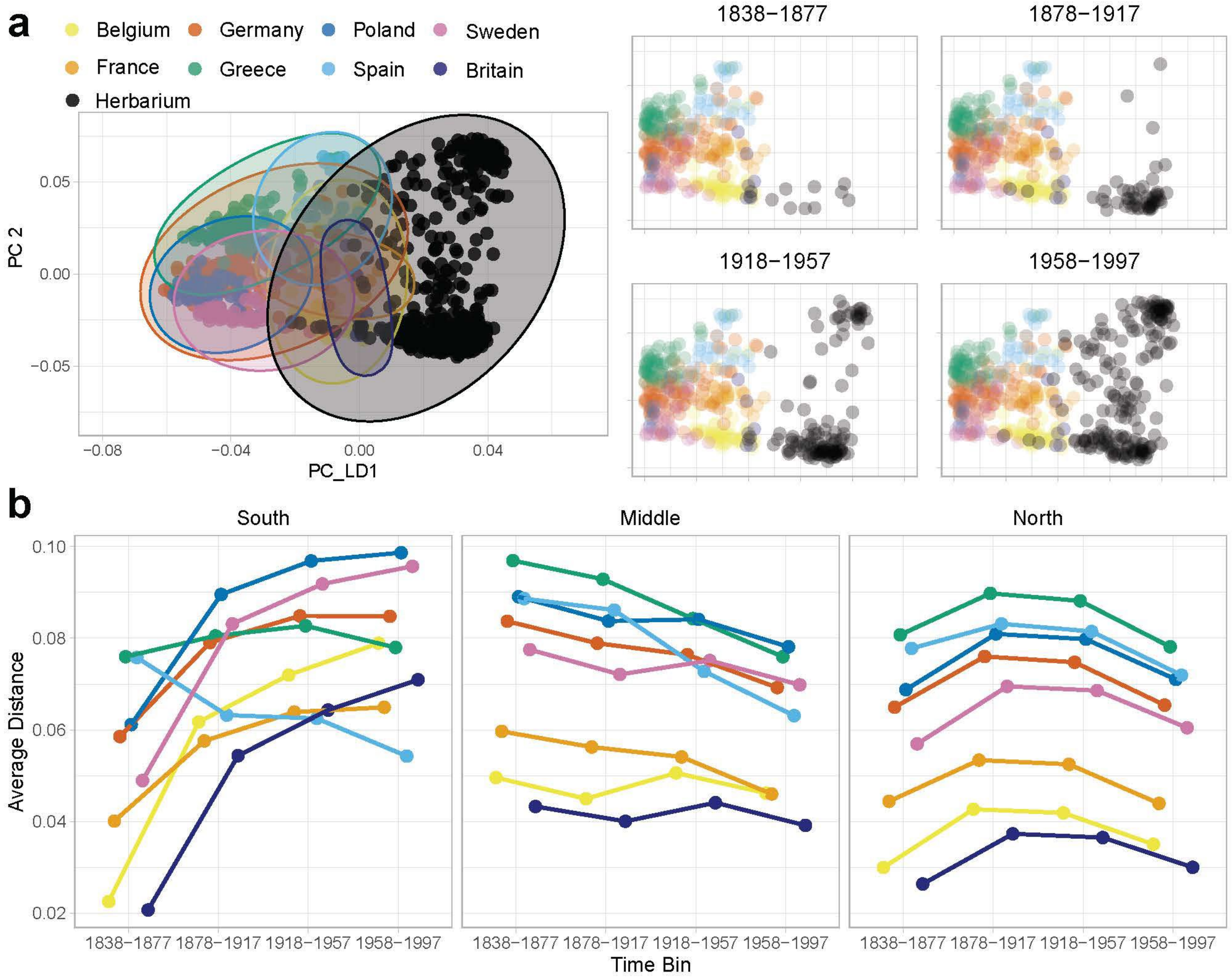

**Fig. S4** TCS haplotype network of chloroplast genomes from herbarium specimens (1838–present) and native European populations (Britain, Spain, and France). Herbarium samples are grouped into 40-year time bins (1838–1877, 1878–1917, 1918–1957, 1958–1997, 1998–present). Each circle represents a unique haplotype, with circle size proportional to the number of samples. Pie charts indicate the temporal or geographic composition of samples within each haplotype (colors correspond to time periods and European source populations). Hash marks along branches denote mutational steps between haplotypes. Only haplotypes represented by ≥2 individuals are shown. The network is based on 307 filtered chloroplast variant sites.

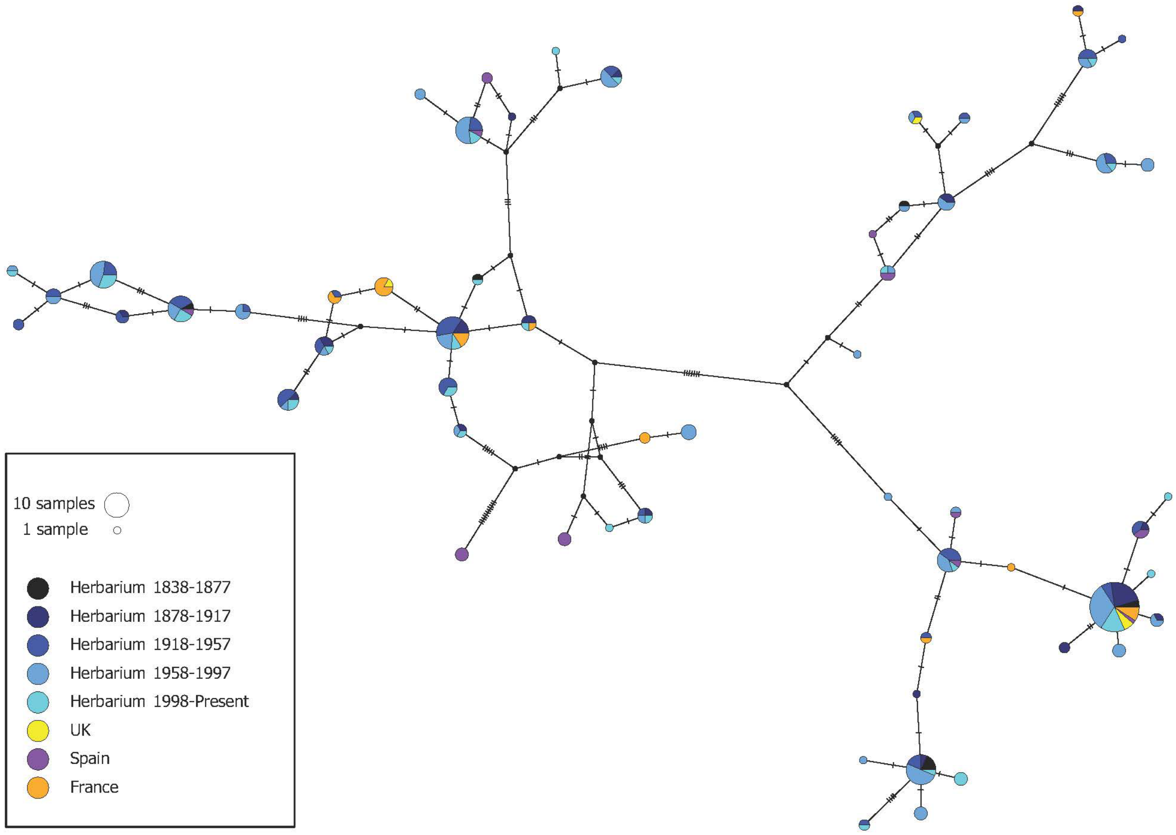

**Fig. S5** Variation in haploblock genotypes over time. Polynomial regressions of haploblock genotypes for 5 structural inversions across time in the north (blue), south (red), and continent wide (black). Points are individual samples. All regression lines are nonsignificant.

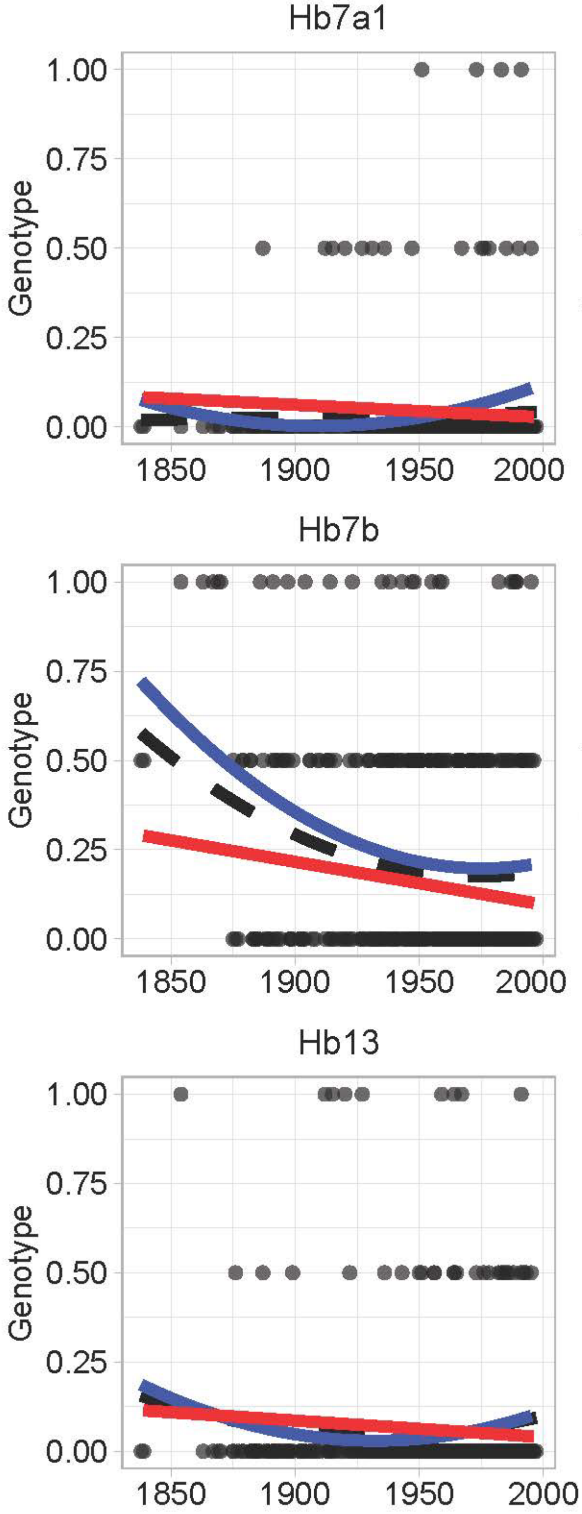

**Table S1** Sample metadata, sequencing quality metrics, population genetic statistics, and genotype assignments for all Trifolium repens herbarium and contemporary specimens.

**Table S2** Genomic coordinates and chromosomal locations of haploblocks identified in *Trifolium repens* used in herbarium specimen analyses. Haploblock regions were identified following Battlay et al. (2025) and extracted from genome-wide genotype likelihood files generated using ANGSD. Principal component analysis was performed on each haploblock region independently using PCAngsd. Table reports the haploblock identifier, chromosome, and start and end coordinates (bp) in the *T. repens* genome assembly.

| Haploblock ID | Chr | Start | End |
|---|---|---|---|
| hb7a1 | Chr04_Occ | 1 | 3720000 |
| hb7a2 | Chr04_Occ | 1400001 | 8500000 |
| hb7b | Chr04_Occ | 50800001 | 54500000 |
| hb9 | Chr05_Occ | 3900001 | 5100000 |
| hb13 | Chr07_Occ | 1 | 1800000 |

**Table S3** Random forest classification results for herbarium specimens assigned to European source populations. The best-performing model was selected based on the lowest out-of-bag (OOB) error rate across all iterations. Confusion matrix from the best-performing model showing classification accuracy for each European source country in the training set, with OOB error rates per class. Summary of predicted country-of-origin assignments for North American herbarium specimens from the best-performing model, including the number of specimens assigned to each source population.

| OOB : 14.58% | | | | | | | | | |
|---|---|---|---|---|---|---|---|---|---|
| **Confusion Matrix** | Belgium | France | Germany | Greece | Poland | Spain | Sweden | UK | class.error |
| Belgium | 5 | 1 | 0 | 0 | 0 | 0 | 0 | 0 | 0.1667 |
| France | 0 | 4 | 0 | 0 | 0 | 0 | 0 | 2 | 0.333 |
| Germany | 0 | 0 | 5 | 0 | 0 | 1 | 0 | 0 | 0.1667 |
| Greece | 0 | 0 | 0 | 6 | 0 | 0 | 0 | 0 | 0 |
| Poland | 0 | 0 | 1 | 0 | 5 | 0 | 0 | 0 | 0.1667 |
| Spain | 0 | 0 | 0 | 0 | 0 | 6 | 0 | 0 | 0 |
| Sweden | 0 | 0 | 0 | 0 | 0 | 0 | 6 | 0 | 0 |
| UK | 0 | 2 | 0 | 0 | 0 | 0 | 0 | 4 | 0.333 |
| **Best Model** | | | | | | | | | |
| prediction | N | Count | sd | se | ci | iteration | | | |
| Belgium | 6 | 1 | 0 | 0 | 0 | 834 | | | |
| France | 34 | 1 | 0 | 0 | 0 | 834 | | | |
| Spain | 143 | 1 | 0 | 0 | 0 | 834 | | | |
| UK | 280 | 1 | 0 | 0 | 0 | 834 | | | |

*Random forest models were trained on the first two principal components (PC1 and PC2) derived from a covariance matrix of genome-wide SNP data to classify North American herbarium specimens to European country of origin (Belgium, France, Spain, Britain, Sweden, Poland, Greece).

*Training sets consisted of European specimens with balanced sampling of up to six individuals per country per iteration to account for unequal sample sizes across source populations.

*Models were run with 5,000 decision trees across 1,000 bootstrap iterations, each with independent random subsampling of European training individuals.

**Table S4** Linear and quadratic model results for temporal trends in NGSadmix cluster proportions across latitudinal regions of North America. Linear and quadratic regression models were fit to examine temporal trends (1838–1997) in maximum cluster assignment (Max Q) and the proportional membership of each specimen to cluster 1 (Q1) and cluster 2 (Q2). Models were run separately for three latitudinal regions: North (>40°N), Mid-Atlantic (35°N–40°N), and South (<35°N).

| Response | Region | Best model | Linear AIC | Quadratic AIC | Linear p-value | Linear Estimate | Linear F-statistic | Quadratic p-value | Quadratic F-statistic |
|---|---|---|---|---|---|---|---|---|---|
| Max Q | North | quadratic | -189.8387 | -213.148 | 0.0005 | -0.0011 | 12.8299 | <0.0001 | 20.9326 |
| Max Q | South | linear | -75.7479 | -73.8032 | 0.7483 | -0.0003 | 0.1036 | 0.819 | 0.0781 |
| Max Q | Mid-Atlantic | linear | -86.4545 | -85.2573 | <0.0001 | -0.0025 | 22.0295 | 0.402 | 11.3812 |
| Cluster Q2 | North | linear | -114.4037 | -112.4362 | 0.1996 | 0.0005 | 1.6591 | 0.8484 | 0.8404 |
| Cluster Q2 | South | linear | 78.0711 | 80.0703 | 0.0099 | 0.0044 | 6.908 | 0.9917 | 3.4199 |
| Cluster Q2 | Mid-Atlantic | quadratic | 41.7574 | 40.5835 | 0.0317 | 0.0022 | 4.7494 | 0.0768 | 3.9903 |
| Cluster Q1 | North | quadratic | -57.3568 | -62.3728 | 0.002 | -0.0014 | 9.8365 | 0.0097 | 8.6263 |
| Cluster Q1 | South | linear | 53.2027 | 54.8481 | 0.0025 | -0.0046 | 9.6494 | 0.5727 | 4.9655 |
| Cluster Q1 | Mid-Atlantic | quadratic | 46.6802 | 45.8857 | 0.0055 | -0.003 | 8.0606 | 0.0954 | 5.4767 |

*Clusters based on NGSadmix estimates of Best K = 3.

*For each response variable and region, both a linear model (response ~ Year) and a quadratic model (response ~ Year + $Year^2$) were fit using ordinary least squares.

*South region analyses were restricted to specimens collected after 1900 given sparse early sampling.

*All models were restricted to specimens collected before 1998.

**Table S5** Linear and quadratic model results for temporal trends in DAPC-derived European ancestry proportions across latitudinal regions of North America.

| Country | Region | Best model | Linear AIC | Quadratic AIC | Linear p-value | Linear Estimate | Linear F-statistic | Quadratic p-value | Quadratic F-statistic |
|---|---|---|---|---|---|---|---|---|---|
| Spain | South | linear | 51.392 | 52.015 | 0.007 | 0.007 | 8.062 | 0.259 | 4.713 |
| Spain | North | linear | -98.665 | -97.331 | 0.207 | 0.0005 | 1.617 | 0.424 | 1.128 |
| Spain | Mid-Atlantic | quadratic | 6.985 | 5.792 | 0.497 | 0.001 | 0.466 | 0.082 | 1.801 |
| Britain | South | linear | -18.577 | -18.508 | 0.013 | -0.003 | 6.647 | 0.181 | 4.308 |
| Britain | North | quadratic | -73.826 | -87.839 | 0.002 | -0.001 | 10.367 | 8.73E-05 | 14.594 |
| Britain | Mid-Atlantic | quadratic | -21.167 | -24.246 | 0.002 | -0.002 | 9.993 | 0.028 | 7.841 |
| France | South | linear | 5.881 | 7.4 | 0.811 | 0.0004 | 0.058 | 0.504 | 0.255 |
| France | North | linear | -118.138 | -116.138 | 0.146 | 0.0005 | 2.15 | 0.996 | 1.063 |
| France | Mid-Atlantic | linear | 0.403 | 2.319 | 0.016 | 0.002 | 6.109 | 0.778 | 3.049 |
| Belgium | South | linear | -19.394 | -17.421 | 0.001 | -0.004 | 13.414 | 0.873 | 6.578 |
| Belgium | North | quadratic | -40.871 | -54.35 | 0.433 | 0.0004 | 0.621 | <0.0001 | 8.527 |
| Belgium | Mid-Atlantic | linear | -3.202 | -1.297 | 0.595 | -0.0005 | 0.285 | 0.765 | 0.186 |

*DAPC estimates were calculated for eight European source populations (Belgium, Spain, United Kingdom, France, Poland, Germany, Sweden, Greece) across years and regions. Displayed analyses are restricted to the populations that exhibited significant temporal changes across introduced regions.

*DAPC was run using ten PCA axes and two discriminant axes (n.pca = 10, n.da = 2), and posterior membership probabilities for each North American specimen were normalized to sum to one across the eight European source populations before averaging within each year.

*Mean annual ancestry proportions were then regressed against year using both linear (Proportion ~ Year) and quadratic (Proportion ~ Year + Year$^2$) ordinary least squares models for each country-by-region combination.

*All analyses were restricted to specimens collected before 1998.

**Table S6** Annual mean DAPC-derived European ancestry proportions for North American herbarium specimens of Trifolium repens by latitudinal region and source country.

**Table S7** Linear model results for the effect of collection year and latitude on genome-wide heterozygosity in North American herbarium specimens of *Trifolium repens*. Ordinary least squares regression models were fit to examine the relationship between collection year and individual-level genome-wide heterozygosity in North American herbarium specimens collected between 1838 and 1997.

| Region | term | Estimate | Std Error | p-value | F-statistic |
|---|---|---|---|---|---|
| All | (Intercept) | -0.04 | 0.017 | 0.02 | 10.31 |
| All | Year | 2.16E-05 | 8.76E-06 | 0.014 | |
| All | Latitude | 0.001 | 0.0004 | 0.029 | |
| All | Year x Latitude | -4.66E-07 | 2.18E-07 | 0.033 | |
| North | (Intercept) | -0.001 | 0.002 | 0.607 | 3.218 |
| North | Year | 2.23E-06 | 1.24E-06 | 0.075 | |
| South | (Intercept) | -0.013 | 0.005 | 0.01 | 10.47 |
| South | Year | 7.84E-06 | 2.42E-06 | 0.002 | |
| Mid-Atlantic | (Intercept) | -1.30E-02 | 4.00E-03 | 0.002 | 15.63 |
| Mid-Atlantic | Year | 8.13E-06 | 2.06E-06 | 0.0002 | |

*A global model (All) including collection year, latitude, and their interaction (Heterozygosity ~ Year × Latitude) was fit across all specimens with valid latitudinal region assignments.

*Sub-analyses were then conducted separately for three latitudinal regions: North (>40°N), Mid-Atlantic (35°N–40°N), and South (<35°N), each fit with collection year as the sole predictor (Heterozygosity ~ Year).

*Model fit was assessed using DHARMa-simulated residuals, including tests for uniformity, dispersion, and outliers.

*All analyses were restricted to specimens collected before 1998.

**Table S8** Linear model results for the effect of maximum NGSadmix cluster assignment (Max Q) and latitude on genome-wide heterozygosity in North American herbarium specimens of *Trifolium repens*. Linear models were fit to examine the relationship between maximum NGSadmix cluster assignment proportion at K=3 (Max Q) and individual-level genome-wide heterozygosity.

| Region | term | Estimate | Std. Error | p-value | F-statistic |
|---|---|---|---|---|---|
| All | (intercept) | 6.91E-03 | 1.20E-03 | 1.65E-08 | 11.69 |
| All | Max Q | 5.44E-03 | 1.40E-03 | 0.000119 | |
| All | Latitude | -9.09E-05 | 3.15E-05 | 0.004099 | |
| All | Max Q x Lat | 1.27E-04 | 3.69E-05 | 0.00063 | |
| North | (intercept) | 3.04E-03 | 2.81E-04 | <2E-16 | 0.02366 |
| North | Max Q | 5.02E-05 | 3.27E-04 | 0.878 | |
| South | (intercept) | 4.25E-03 | 2.96E-04 | <2E-16 | 22.39 |
| South | Max Q | -1.61E-03 | 3.41E-04 | 7.26E-06 | |
| Mid-Atlantic | (intercept) | 3.41E-03 | 3.04E-04 | <2E-16 | 3.723 |
| Mid-Atlantic | Max Q | -7.11E-04 | 3.68E-04 | 0.0566 | |

*A global model (All) including Max Q, latitude, and their interaction (Heterozygosity ~ Max Q × Latitude) was fit across all specimens with valid latitudinal region assignments.

*Sub-analyses were then conducted separately for three latitudinal regions: North (>40°N), Mid-Atlantic (35°N–40°N), and South (<35°N), each fit with Max Q as the sole predictor (Heterozygosity ~ Max Q).

*Model fit was assessed using DHARMa-simulated residuals, including tests for uniformity, dispersion, and outliers.

*All analyses were restricted to specimens collected before 1998.

**Table S9** Linear model results for the effect of collection year and latitude on standardized *Ac* and *Li* locus frequencies in North American herbarium specimens of *Trifolium repens*. Ordinary least squares regression models were fit to examine temporal trends in standardized cyanogenesis locus frequencies (*Ac/ac* and *Li/li*) in North American herbarium specimens collected between 1838 and 1997.

| stdAC | Region | term | estimate | Std Error | p-value | F-statistic |
|---|---|---|---|---|---|---|
| | All | (Intercept) | -7.33E-05 | 2.68E-05 | 0.006 | 3.814 |
| | All | Year | 3.83E-08 | 1.36E-08 | 0.005 | |
| | All | Latitude | 1.83E-06 | 6.71E-07 | 0.007 | |
| | All | Year x Latitude | -9.41E-10 | 3.43E-10 | 0.006 | |
| | North | (Intercept) | 2.87E-06 | 4.33E-06 | 0.508 | 0.1746 |
| | North | Year | -9.32E-10 | 2.23E-09 | 0.677 | |
| | South | (Intercept) | -7.39E-06 | 7.09E-06 | 0.3 | 1.5 |
| | South | Year | 4.42E-09 | 3.61E-09 | 0.224 | |
| **stdLI** | **Region** | **term** | **Estimate** | **Std Error** | **p-value** | **F-statistic** |
| | All | (Intercept) | -0.00014 | 4.23E-05 | 0.001 | 43.37 |
| | All | Year | 7.41E-08 | 2.16E-08 | 0.001 | |
| | All | Latitude | 3.19E-06 | 1.06E-06 | 0.003 | |
| | All | Year x Latitude | -1.70E-09 | 5.42E-10 | 0.002 | |
| | North | (Intercept) | -3.15E-06 | 5.02E-06 | 0.531 | 0.7338 |
| | North | Year | 2.22E-09 | 2.59E-09 | 0.393 | |
| | South | (Intercept) | -3.99E-05 | 1.44E-05 | 0.007 | 8.95 |
| | South | Year | 2.20E-08 | 7.34E-09 | 0.003 | |

*Global models (All) including collection year, latitude, and their interaction (Frequency ~ Year × Latitude) were fit across all specimens to test for latitudinally divergent temporal trends.

*Sub-analyses were then conducted separately for three latitudinal regions: North (>40°N), Mid-Atlantic (35°N–40°N), and South (<35°N), each fit with collection year as the sole predictor (Frequency ~ Year).

*Analyses were restricted to specimens collected before 1998.

**Table S10** Linear model results for the effect of collection year and latitude on haploblock frequency in North American herbarium specimens of *Trifolium repens*. Ordinary least squares regression models were fit to examine temporal trends in haploblock frequencies across North American herbarium specimens collected between 1838 and 1997.

| Haploblock | Region | Best model | Linear AIC | Quadratic AIC | Linear p-value | Quadratic p-value |
|---|---|---|---|---|---|---|
| hb7a1 | All | linear | -386.49 | -385.38 | 0.134 | 0.351 |
| hb7a1 | North | quadratic | -126.35 | -126.83 | 0.585 | 0.122 |
| hb7a1 | South | linear | -88.46 | -86.61 | 0.715 | 0.7 |
| hb7a2 | All | linear | -72.55 | -70.97 | 0.215 | 0.52 |
| hb7a2 | North | quadratic | -40.03 | -41.79 | 0.107 | 0.057 |
| hb7a2 | South | linear | -7.56 | -5.69 | 0.027 | 0.725 |
| hb7b | All | quadratic | 162.47 | 161.33 | 0.506 | 0.079 |
| hb7b | North | quadratic | 100.3 | 100.11 | 0.499 | 0.145 |
| hb7b | South | linear | 23.37 | 24.38 | 0.187 | 0.334 |
| hb9 | All | quadratic | -232.6 | -233.79 | 0.193 | 0.076 |
| hb9 | North | linear | -131.23 | -131.04 | 0.729 | 0.186 |
| hb9 | South | linear | -65.64 | -64.18 | 0.294 | 0.475 |
| hb13 | All | quadratic | -148 | -148.32 | 0.529 | 0.131 |
| hb13 | North | quadratic | -71.26 | -71.87 | 0.023 | 0.112 |
| hb13 | South | linear | -50.54 | -48.78 | 0.09 | 0.633 |

*Global models (All) including collection year, latitude, and their interaction (Haploblock Frequency ~ Year × Latitude) were fit across all specimens.

*Sub-analyses were conducted separately for three latitudinal regions: North (>40°N), Mid-Atlantic (35°N–40°N), and South (<35°N), each fit with collection year as the sole predictor.

*All analyses were restricted to specimens collected before 1998.
